# The *Arabidopsis* TNL immune receptor BNT1 localizes to the plastid envelope and mediates flg22-induced resistance against *Pseudomonas*

**DOI:** 10.1101/2025.01.29.635455

**Authors:** Micaela Y. Peppino Margutti, Ana P. Cislaghi, Ariel Herrera-Vásquez, Julieta R. Palomeque, Francisco Bellino, María E. Alvarez, Francisca Blanco-Herrera, Nicolás M. Cecchini

## Abstract

Precise localization and trafficking of plant immune receptors are critical for their function. We identify the TNL-class nucleotide-binding leucine-rich repeat receptor (NLR) BURNOUT1 (BNT1) from *Arabidopsis thaliana* as localized to plastids, key organelles for plant immunity. Alternative transcription start site usage generates two isoforms of BNT1: BNT1.2, which is targeted to the plastid envelope via an N-terminal signal-anchored mechanism, and BNT1.1, which resides in the cytoplasm. Moreover, BNT1.2 is predominantly expressed in epidermal cells, where it localizes to the so-called sensory plastids. Functional analysis revealed that *bnt1* mutants exhibit compromised PAMP-triggered immunity (PTI) responses, including impaired callose deposition and reduced flg22-induced resistance to *Pseudomonas syringae* pv. *tomato*, while flg22-induced apoplastic reactive oxygen species production remains unaffected. Notably, only the plastid-localized BNT1.2 isoform is required for these PTI responses. Our findings reveal a role for NLRs in regulating PTI responses from plastids and highlight these organelles as key hubs for signal(s) integration during plant-pathogen interactions.

**Significance statement:** This study identifies BNT1 as a TNL-class immune receptor localized to the plastid envelope. Two distinct isoforms of BNT1 were characterized: one with a plastid-targeting signal anchor that ensures plastid localization and another confined to the cytoplasm. Notably, only the plastid-localized isoform mediates PTI responses and confers resistance to *Pseudomonas*, highlighting the critical role of precise NLR localization and the central role of plastids in plant immunity.

## INTRODUCTION

The ability of plant immune system to recognize pathogens depends mostly on two types of proteins, the <u>p</u>attern recognition receptors (PRRs) that reside on the cell surface and perceive pathogen-associated molecular patterns (PAMPs; *e.g.* flg22 peptide from bacterial flagellin) (Macho and Zipfel, 2014), and the intracellular resistance <u>n</u>ucleotide-binding leucine-rich repeat receptors (NLRs) that sense specific effector proteins used by microbes to promote their virulence (Jones *et al*., 2016). Activation of PRRs or NLRs induces the so-called PAMP-triggered immunity (PTI) or effector-triggered immunity (ETI), respectively (Jones *et al*., 2024). PTI and ETI act synergistically, boosting each other to confer full immunity (Ngou *et al*., 2021; Bernoux *et al*., 2022; Chen *et al*., 2022; Yu *et al*., 2024). These pathways share several downstream defense responses and signaling mechanisms, such as activation of mitogen-activated protein kinase (MAPK) cascades, intracellular calcium increases, production of reactive oxygen species (ROS), deposition of callose for cell wall reinforcement (Weralupitiya *et al*., 2024; Dodds *et al*., 2024) and transcriptional reprogramming, including the induction of defense genes (*e.g. PATHOGENESIS-RELATED GENE 1* (*PR1*) (Li *et al*., 2001; Tsuda *et al*., 2009). Furthermore, the activation of both PRRs and/or NLRs is known to trigger a state of enhanced defenses against future pathogen attacks, known as induced resistance (IR) (Zipfel *et al*., 2004; Mishina and Zeier, 2007; De Kesel *et al*., 2021; Hönig *et al*., 2023).

NLRs recognize the pathogen effectors either directly by binding to them, or indirectly, by sensing alterations in their virulence targets or target decoys, triggering robust defense responses (Jones *et al*., 2016; Cesari, 2018; Contreras *et al*., 2023). Therefore, maintaining an extensive and balanced repertoire of NLRs is crucial for an effective and controlled plant immune system. NLRs diversity and abundance are regulated by multiple mechanisms, including transcript processing, such as alternative transcription start sites (TSS) and alternative splicing (Li *et al*., 2015; Kourelis and Adachi, 2022; Shepherd *et al*., 2023; Sun *et al*., 2024). The structure of NLR proteins consist of a conserved nucleotide-binding domain (NB), a leucine-rich repeat (LRR) domain and a variable N-terminal region that can resemble either the “Toll/Interleukin-1 receptor” (TIR) or a “coiled coil” (CC) domain, and thus are referred as TIR/CC-NB-LRR (or TNL/CNL) (Kourelis *et al*., 2021a). Moreover, CNL subclasses exist that contain variants of the CC domain, such as the so-called helper NLRs, which have a CC domain homologous to that of the *Resistance to Powdery Mildew 8* (RPW8) protein family (RPW8-like; CC_R_-NLRs) (Xiao *et al*., 2004; Kourelis *et al*., 2021b). The helper CC_R_-NLRs assist in the signaling pathway downstream of other NLR receptors, without being directly involved in effectors’ perception (Jubic *et al*., 2019; Ngou *et al*., 2020; Gong *et al*., 2023). TNLs and CNLs can function as monomers, self-associate, assemble into hetero-or homodimers, or form higher-order oligomers, such as the wheel-shaped “resistosomes” (Cesari, 2018; Adachi *et al*., 2019; Tamborski and Krasileva, 2020), with some CNL-resistosomes forming plasma membrane pores to trigger calcium influx and defense responses (Huang *et al*., 2023; Shepherd *et al*., 2023).

NLRs are distributed across various subcellular compartments like the nucleus, plasma membrane, cytoplasm, and endomembrane system, and their precise location and trafficking appear to be crucial for their function (Padmanabhan and Dinesh-Kumar, 2010; Qi and Innes, 2013; Duggan *et al*., 2021; Lüdke *et al*., 2022; Bernoux *et al*., 2023; Shepherd *et al*., 2023). Moreover, a single CC_R_-NLR was recently located in the chloroplast, although the significance of its localization for plant immunity remains unclear (Ibrahim *et al*., 2024). Plastids are directly related to numerous processes associated with PTI, ETI, and IR programs (Caplan *et al*., 2008; Nomura *et al*., 2012; Zabala *et al*., 2015; Serrano *et al*., 2016; Medina-Puche *et al*., 2020; Kachroo *et al*., 2021; Littlejohn *et al*., 2021; Yang *et al*., 2021). They produce key defense phytohormones and defense-signals, generate reactive oxygen species (ROS), and may serve as a calcium source (Grant and Jones, 2009;Jung *et al*., 2009; Zurbriggen *et al*., 2009; Nomura *et al*., 2012; Shapiguzov *et al*., 2012; Zeier, 2013; Cecchini et al. 2015a; Stael *et al*., 2015; Mignolet-Spruyt *et al*., 2016; Serrano *et al*., 2016; Salinas et. al 2024; Weralupitiya *et al*., 2024). During PTI and ETI, plastids move and cluster around the nucleus, forming stromules that are proposed to facilitate retrograde signaling (Caplan *et al*., 2015; Noctor and Foyer, 2016; Park *et al*., 2018; Ding *et al*., 2019; Hanson and Conklin, 2020; Jung *et al*., 2024). Moreover, plastid components are targeted by many effectors from bacteria, oomycetes, fungi, and sucking insects (Jelenska *et al*., 2007; Li *et al*., 2014; Zabala *et al*., 2015; Mugford *et al*., 2016; Rosas-Diaz *et al*., 2018; Kretschmer *et al*., 2019; Lee and Hwang, 2020; Medina-Puche *et al*., 2020; Tzelepis *et al*., 2021; Yang *et al*., 2021; Lovelace, 2023; Stojilković *et al*., 2024). Thus, while plastids play essential roles in immune signaling, the involvement of plastid-targeted NLRs in plant immunity remains largely unexplored.

We recently characterized an N-terminal bipartite signal for plastid localization, which was thereafter used to predict plastid-targeted proteins in *Arabidopsis thaliana* (hereafter *Arabidopsis*) (Cecchini *et al*., 2021; Banday *et al*., 2022). Remarkably, this led to the identification of several NLRs harboring this signal (Banday *et al*., 2022). Here, we show that one of them, the TNL-class BURNOUT 1 (AT5G11250/BNT1; (Sarazin *et al*., 2015)), is located at the plastids envelope. We report two BNT1 isoforms generated through the use of alternative transcription start sites (TSS); one includes an extended exon encoding the plastid-targeting signal, while the other localizes in the cytoplasm. Moreover, the characterization of *bnt1* mutant plants revealed that the plastid-localized BNT1 isoform is required for PTI responses, including callose deposition and flg22-induced resistance (flg22-IR) against *Pseudomonas syringae* pv. *tomato*. Our study highlights the importance of NLR subcellular localization for their function and underscores plastids as key organelles in plant immunity.

## RESULTS

### BNT1 is targeted to the plastid envelope

We recently identified a predicted chloroplast localization signal in the *Arabidopsis* TNL-class NLR AT5G11250.2/BNT1.2 isoform (Banday *et al*., 2022). Supporting this, BNT1 peptides were found in a proteomic study of plastids (Beltrán *et al*., 2018). Thus, we generated a construct with the full-length *BNT1.2* (amino acids 1 to 1189) fused to the green fluorescent protein under dexamethasone (Dex)-inducible promoter (BNT1.2-GFP) and analyzed its subcellular localization.

Confocal microscopy of *Nicotiana benthamiana* leaves transiently expressing BNT1.2-GFP showed GFP signal at plastids, exhibiting a ring-like pattern characteristic of outer envelope membrane proteins, as well as in stromules of varying lengths (Figure 1a; Movie S1). This finding was further validated through subcellular fractionation. BNT1.2-GFP was highly enriched in the plastid fraction compared to the endoplasmic reticulum (ER)-anchored defense protein TN13 (Figure 1b) (Banday *et al*., 2022). To better characterize BNT1 localization, we co-expressed BNT1.2-GFP with the outer envelope membrane marker OEP7 or the ER marker BiP tagged with red fluorescence protein (RFP) (OEP7-RFP or BiP-RFP) (Cecchini *et al*., 2015b). The microscopy analysis revealed that most of the BNT1.2-GFP signal overlapped with OEP7-RFP. In contrast, a smaller portion co-localized with BiP-RFP, particularly at contact sites between plastid stromules and the ER (Figure 1c,d). The fluorescence intensity profiling of the merged images further supported this observation (Figure 1d,e right graphs). Next, we generated BNT1.2-GFP *Arabidopsis* transgenic plants and studied BNT1 location. As shown in Figure 1e, the plastid envelope localization pattern of BNT1.2-GFP closely resembled the one observed in *N. benthamiana*. Interestingly, although expressed under a non-tissue specific Dex-inducible promoter, BNT1.2-GFP plastid signal was only detected in *Arabidopsis* leaf epidermal cells (Figure 1e and Figure S1; Movie S2).

**Figure 1.**
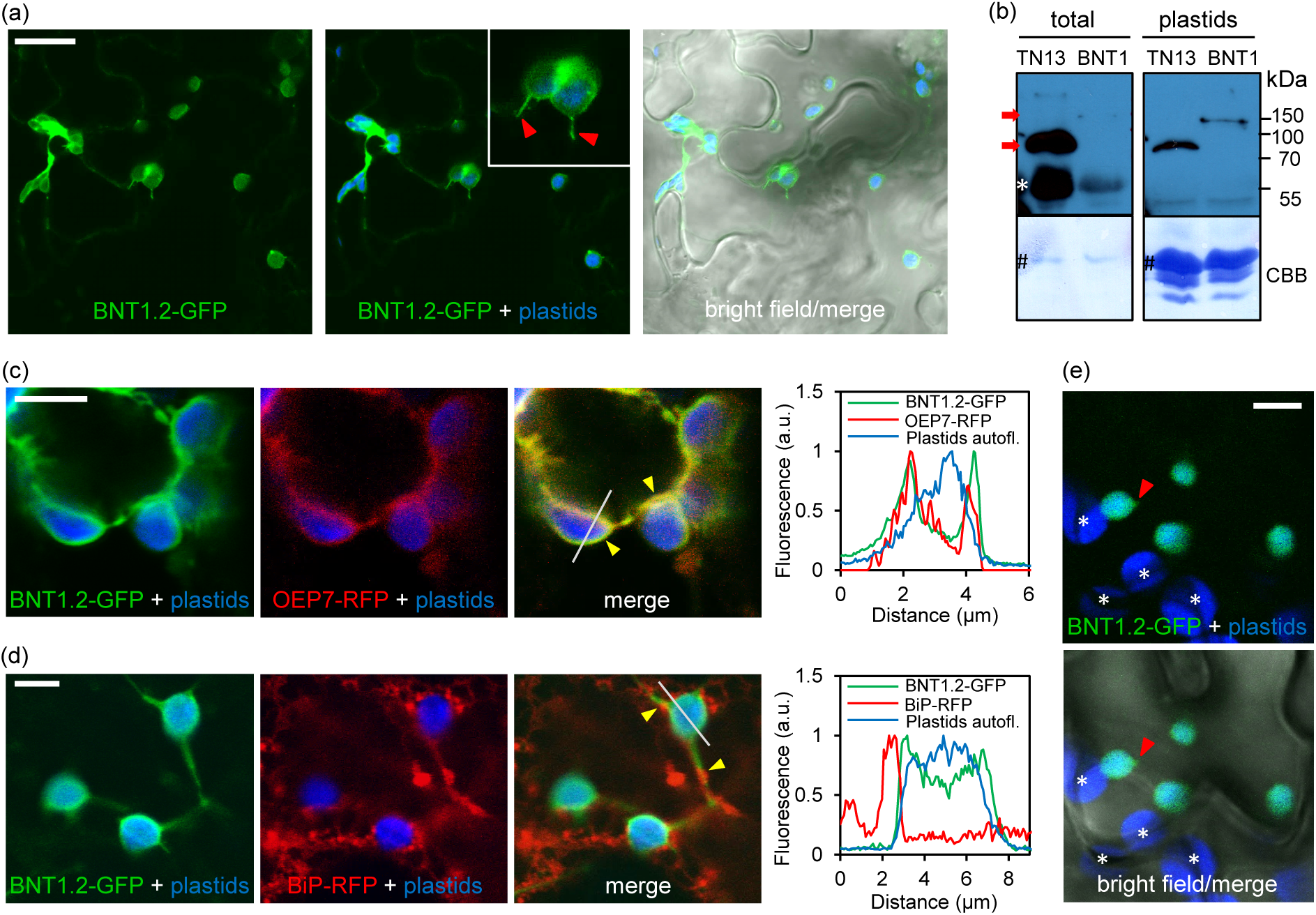
Subcellular localization of BNT1. (a) Laser scanning confocal microscopy (confocal) micrographs showing localization of GFP-tagged BNT1.2 controlled by Dex-in-ducible promoter (BNT1.2-GFP) transiently expressed in *Nicotiana benthamiana* epidermal cells. Red arrowheads: stromules. (b) Western blots of total and plastid fractions from N. benthamiana leaves expressing GFP-tagged TN13 or BNT1.2. Bands were revealed using anti-GFP antibody. The blots stained with Coomassie blue (CBB) are presented to show loading and Rubisco (#). Red arrows indicate expected band sizes. Asterisk show unspecific bands. (c,d) Confocal micrographs showing BNT1.2-GFP co-expressed in *N. benthamiana* with a plastid outer envelope protein marker (OEP7:RFP) (c) or ER marker (BiP:RFP) (d). Yellow arrowheads: co-localization. Right graphs: fluorescence profiles for green and red channels along the white lines shown in the merge micrographs. a.u.: arbitrary units. (e) Confocal micrographs showing BNT1.2-GFP in leaves of transgenic *Arabidopsis thaliana*. Red arrowheads: stromule. Asteris-ks indicate spongy mesophyll cells plastids. Micrographs show GFP (green), RFP (red) and plastid autofluorescence (blue). In (b) Bar = 20 μm, and (d-f) Bar = 5 μm.

These results indicate that the predicted plastid signal is functional in this TNL-class NLR, as previously observed for other plant defense factors (Cecchini *et al*., 2015b; Banday *et al*., 2022), directing BNT1.2 to the envelope of plastids, particularly in epidermal cells.

### BNT1 is a plastid signal-anchored protein

The N-terminal bipartite signal predicted in BNT1.2 is a variant of the plastid-targeted signal-anchored (SA) proteins (Kim and Hwang, 2013; Cecchini *et al*., 2021; Banday *et al*., 2022). It consists of a signal peptide-like region (SP-like; amino acids 1–23), a positively charged region (CPR; amino acids 24–30), and a predicted chloroplast transit peptide-like region (cTP-like; amino acids 31–56) (Figure 2a). The CPR functions to prevent the cleavage of the SP-like, thereby constituting a transmembrane (TM) anchor domain.

**Figure 2.**
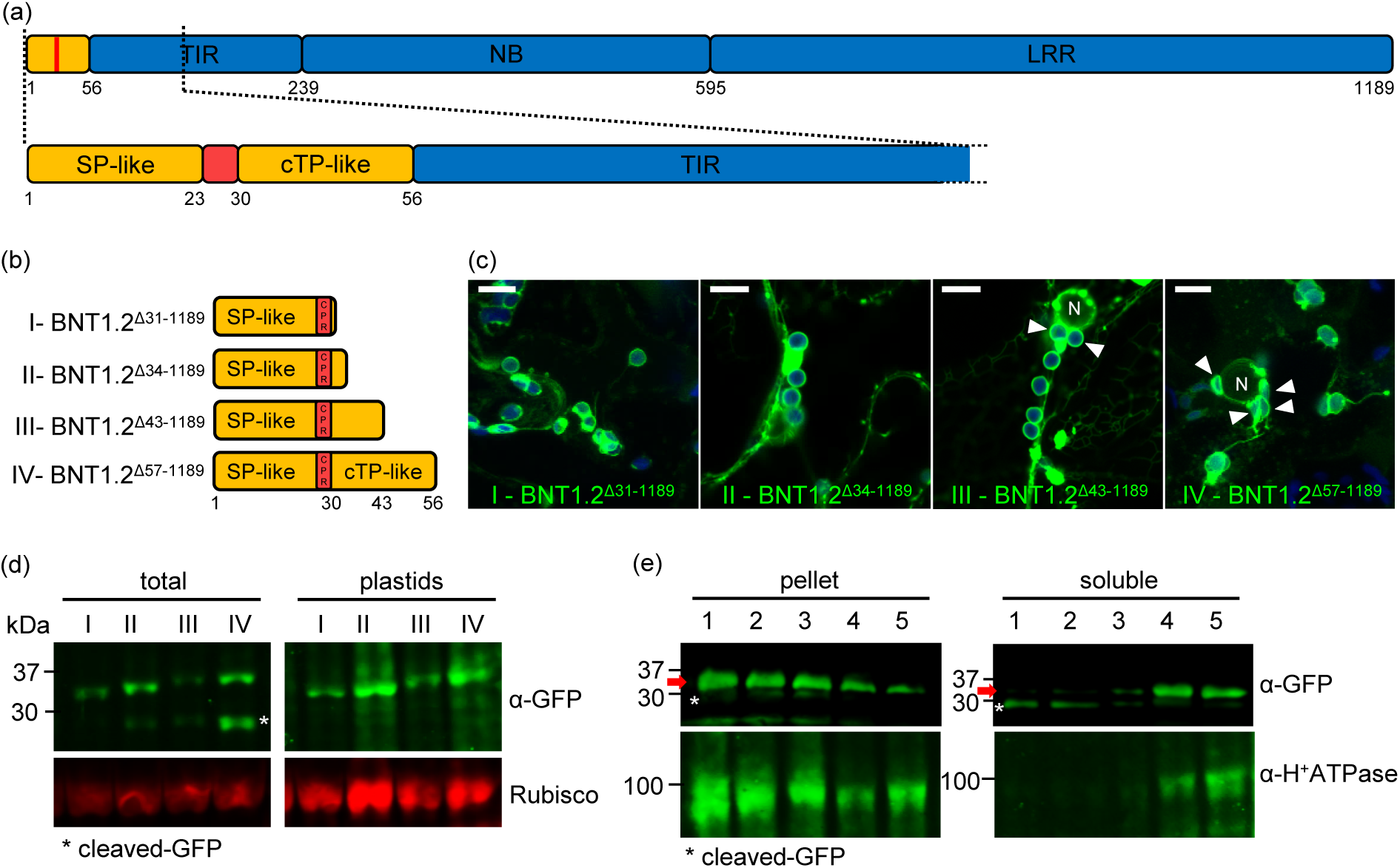
BNT1 N-terminal region is required for plastid targeting and membrane association. (a) Scheme of BNT1.2 showing the N-terminal bipartite signal (yellow, red) and the NLR domains (blue; UniprotKB ID: Q9LFN1_A-RATH). SP-like: predicted signal peptide; cTP-like: predicted chloroplast transit peptide; CPR: positively charged region; interdo-main region (white box), TIR: toll/interleukin-1 receptor domain; NB: nucleotide-binding domain; LRR: leucine-rich repeat. Amino acid positions delimiting regions/domains are shown in the lower part. (b) Schematic of BNT1.2 N-terminal signal deletion variants used for the localization analysis in (c) and fractionations in (d,e). Amino acid positions delimiting regions/domains are shown in the lower part. (c) Z-series maximum intensity projection showing the localization of GFP-tagged BNT1.2N-terminal signal variants (b) in *N. benthamiana*. Bar = 10 µm. White arrowheads: plastids clustered around nucleus (N). (d) Western blots of total or plastid fraction protein from *N. benthamiana* leaves expressing BNT1.2 N-terminal signal variants fused to GFP used in (c). Bands were revealed using anti-GFP antibody. Odyssey 700 nm channel is presented to show loading and Rubisco. (e) Membrane association strength of BNT1.2^Δ31-1189^-GFP variant (b, I-). Western blots of pelleted and soluble proteins after total microsomal fraction treatments with control (150 mM NaCl) (1), 1.5 M NaCl (2), 2 M urea (3); 1% Triton (4), 1% NP-40 + 0.5% deoxycolate (5). Bands were revealed using the indicated antibodies. H+ATPase is an integral membrane protein marker. In (c,d) similar results were observed in two independent experiments. * Indica tes cleaved GFP in (d) and (e).

To effectively evaluate the functionality of the bipartite signal, we generated GFP-tagged deletion variants of the N-terminal of BNT1.2 (Figure 2b) and examined their subcellular localization. Each construct was expressed in *N. benthamiana* leaves and used to perform confocal microscopy and subcellular fractionation analysis. As expected, the complete N-terminal bipartite signal localizes to the plastid envelope (Figure 2c, IV-BNT1.2^Δ57-1189^-GFP), like the full-length BNT1.2 (Figure 1c). However, the analysis showed that the signal-truncated variants exhibited plastid-envelope targeting, including the SP-like+CPR shortest construct (Figure 2c; I, II, III). Fractionation analysis confirmed the enrichment of all GFP fusion constructs in plastid fractions, while free cleaved-GFP was not detected in this fraction (Figure 2d). Interestingly, we noticed that the expression of the complete (or partially complete) bipartite signal constructs (III and IV) appeared to promote clustering of plastids around the nucleus, compared to the other variants.

If BNT1.2 is a SA protein, the SP-like domain will function as a transmembrane (TM) region, anchoring the protein to the membrane. Therefore, we examined the strength and type of membrane association of the shortest BNT1.2 deletion version, the SP-like+CPR (Figure 2b; I-BNT1.2^Δ31-1189^). For this, we isolated the microsomal fraction from *N. benthamiana* leaves expressing BNT1.2^Δ31-1189^-GFP and treated it with different salt concentrations, urea, or detergents. As shown in Figure 2e, only treatments with 1% Triton or 1% NP-40 + 0.5% deoxycolate were effective in extracting BNT1.2^Δ31-1189^-GFP from the microsomal fraction. These same treatments also solubilized the H^+^ATPase, a control marker for membrane proteins. This finding provides substantial evidence that BNT1.2 is a membrane anchored protein.

Together, our data indicate that the complete bipartite signal is not essential for the plastid localization of BNT1.2, suggesting that it utilizes the ‘classic’ SA protein mechanism for targeting to the plastid envelope (Lee *et al*., 2011). The potential role of the complete signal in promoting plastid clustering near nuclei remains to be elucidated.

### BNT1 generates isoforms differing on the plastid-targeting signal

The *Arabidopsis* genome annotation (Araport11, https://www.araport.org/data/araport11) predicts three *BNT1* coding transcript isoforms: *BNT1.1*, *BNT1.2* and *BNT1.3*. Interestingly, *BNT1.1* and *BNT1.2* are generated by alternative transcription start sites (TSSs) that alter their 5′ untranslated region (5’UTR) length (Figure 3a and Figure S2) (CAGE-seq and PEAT-clusters; (Morton *et al*., 2014; Thieffry *et al*., 2020)). The *BNT1.2* transcript contains an extended exon that encodes the N-terminal signal for plastid localization, whereas the *BNT1.1* transcript lacks this signal and the first 20 amino acids of the TIR domain (Figure 3b). In contrast, no alternative TSS was identified in *BNT1.3*, which encodes the same protein as *BNT1.1* (Figure S2), suggesting that it is not a real transcript variant. Supporting this, only *BNT1.1* and *BNT1.2* were detected in a long-read RNA-seq study (Dr. Ezequiel Petrillo, personal communication). To confirm the presence of *BNT1.1* and *BNT1.2* we tested their expression by reverse transcription (RT) followed by semi-quantitative PCR (RT-sqPCR) in *Arabidopsis* leaves. We used specific primers for each variant 5’UTR (Figure 3a). We successfully amplified the transcripts and found that *BNT1.2* is significantly more abundant than *BNT1.1* under basal conditions (Figure 3c). This finding suggests that the isoforms are present and may be subject to differential regulation. As anticipated, the RT-sqPCR negative controls (minus RT) showed no amplification (Figure S3a).

**Figure 3.**
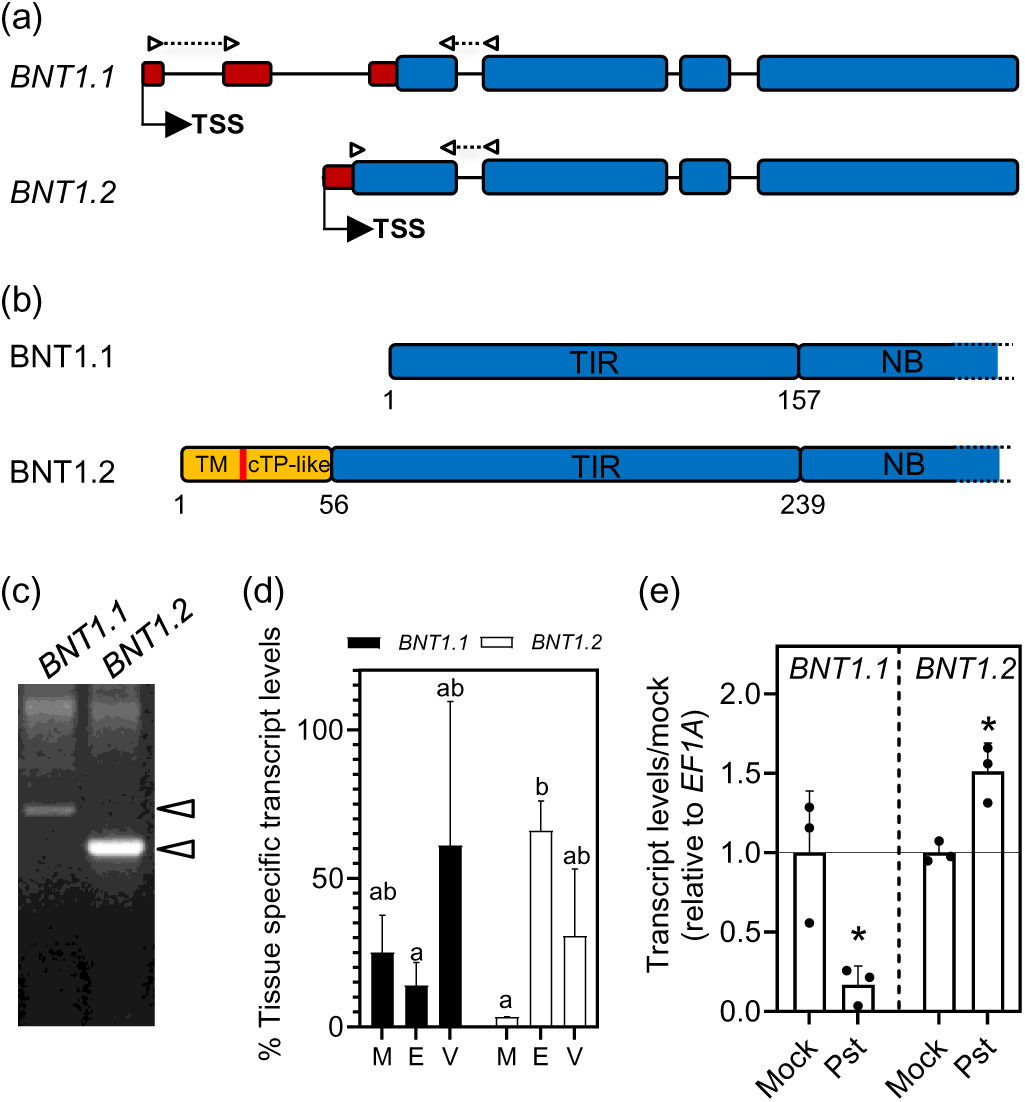
Alternative *BNT1* transcripts encode for isoforms with or without the N-terminal plastid targeting signal and show differential expression. (a) Schematic representation of *BNT1.1* and *BNT1.2* transcripts generated by alternative transcription start sites (TSS). Blue blocks: exons, black lines: introns, red blocks: 5’UTR exons. Arrowheads indicate the position of primers used in (d-e) (dashed lines connecting arrowheads represent genomic sequences). (b) Scheme of BNT1.1 and BNT1.2 isoforms showing protein N-terminal regions/domains. Orange block: N-terminal bipartite targeting signal; red line: CPR region; blue blocks: NLR domains “Toll/Interleukin-1 receptor” (TIR), and part of the nucleotide binding domain (NB). Amino acid positions are indicated with numbers. (c) RT-PCR analysis showing the presence of *BNT1.1* and *BNT1.2* transcripts in *Arabidopsis* adult plant leaves. Arrowheads indicate the expected band sizes. (d) Percentage of *BNT1.1* and *BNT1.2* transcript levels in different leaf tissues under basal conditions (M: mesophyll; E: epidermis; V: vascular bundle). The average from three independent experiments is shown. Different letters indicate statistically significant differences (p < 0.05, analysis of variance (ANOVA), Fisher’s LSD test). Absolute transcript levels of both variants are shown in Figure S3. (e) Transcript levels of *BNT1.1* and *BNT1.2* quantified by RT-qPCR in *Arabidopsis* adult plant leaves after 24h hours post-inoculation (hpi) with *Pseudomonas syringae* pv. *tomato* DC3000 (*Pst*) (OD=0.005) or mock treatment (10 mM MgCl_2_). The mean ± SE from three independent experiments is shown (each data point with 3 leaves from 3 different plants pooled together for RNA extraction). Asterisks indicate statistically significant differences between infected and mock-treated leaves (p < 0.05, Student’s *t* test).

Given that NLR expression is typically responsive to environmental cues (Yang *et al*., 2021) and BNT1 protein has been identified by proteomic studies in epidermal and vascular tissues (Beltrán *et al*., 2018), we explored whether *BNT1* expression varies across plant tissues and under biotic stress conditions by analyzing public expression data using Genevestigator® (Hruz *et al*., 2008). This analysis showed that total *BNT1* levels were higher in leaves than in roots, particularly in veins and midrib tissues (Figure S4a). Furthermore, *BNT1* expression was significantly induced by different biotic stresses, including treatments with PAMPs and infections with virulent and non-virulent *Pseudomonas* strains (Figure S4b). Considering this, we next examined the relative transcript levels of *BNT1.1* and *BNT1.2* in different leaf tissues and in response to *Pseudomonas syringae* pv. *tomato* DC3000 (*Pst*) infection, using reverse transcription followed by quantitative PCR (RT-qPCR) using specific primers (Figure 3a). To analyze tissue-specific expression, we first isolated mesophyll cells, epidermal cells and vascular bundles from *Arabidopsis* leaves (Figure S3b) (Endo *et al*., 2016). Interestingly, the expression analysis showed that *BNT1.2* was significantly enriched in epidermal cells compared to *BNT1.1* transcript (Figure 3d and Figure S3c). Although variable and not statistically significant, both isoforms also appear to show higher expression levels in the vasculature. Next, *Arabidopsis* plants were infected with *Pst* to examine isoform-specific responses. After 24 hours post-infection (hpi), leaves infiltrated with *Pst* (OD_600_=0.005) or mock-treated (10 mM MgCl_2_) were collected, and isoforms abundance was analyzed by RT-qPCR. Interestingly, *BNT1.2* levels were significantly increased relative to the mock treatment, while *BNT1.1* levels were repressed, suggesting antagonistic regulation of these isoforms during *Pst* infection (Figure 3e).

Together, these findings suggest that *BNT1* expression is regulated in an isoform-specific manner, likely through alternative TSS usage, particularly during plant-pathogen interactions and across different leaf tissues.

### BNT1 isoforms are targeted to different subcellular locations

*BNT1.2* generates an isoform located in the plastid envelope (Figures 1 and 3), while the *BNT1.1* transcript is predicted to produce a variant lacking the N-terminal plastid-targeting signal. Despite low transcript abundance, a recent super-resolution Ribo-seq study confirmed ribosomal loading of the *BNT1.1* (Wu *et al*., 2024), supporting its translation. Therefore, we next compared the subcellular localization of each isoform.

Full-length BNT1.1-GFP and BNT1.2-GFP constructs were generated under the constitutive 35S promoter, and their expression was analyzed using confocal microscopy. As previously observed with the Dex-inducible construct (Figure 1a), transient expression of BNT1.2-GFP in *N. benthamiana* localized to the plastid envelope (Figure 4a). In contrast, BNT1.1-GFP showed a cytoplasmic distribution near the plasma membrane, consistent with the absence of the plastid-targeting signal. However, no GFP signal was detected in transgenic *Arabidopsis* lines expressing the 35S constructs (not shown), suggesting post-translational regulation or silencing resulting in very low protein levels. Thus, we generated an *Arabidopsis* transgenic plant carrying BNT1.1-GFP under a Dex-inducible promoter and compared it with the Dex-inducible BNT1.2-GFP line (Figure 1e). Although the GFP signal intensity was low for both isoforms, BNT1.2-GFP displayed a ring-like pattern around plastids, while BNT1.1-GFP exhibited a cytoplasmic distribution (Figure 4b). Furthermore, truncated versions of BNT1.1 and BNT1.2 under 35S promoter were constructed, retaining only the TIR or TM+cTP+TIR domains (BNT1.1^Δ158-1107^-GFP and BNT1.2^Δ240-1189^-GFP, respectively). These versions showed localization patterns analogous to the full-length constructs in both *N. benthamiana* and transgenic *Arabidopsis* plants (Figure 4c,d). Interestingly, these N-terminal truncated variants were expressed at significantly higher levels, enabling a precise visualization in *Arabidopsis*, including the BNT1.2^Δ240-1189^-GFP locating in stromules (Figure 4d, bottom panels).

**Figure 4.**
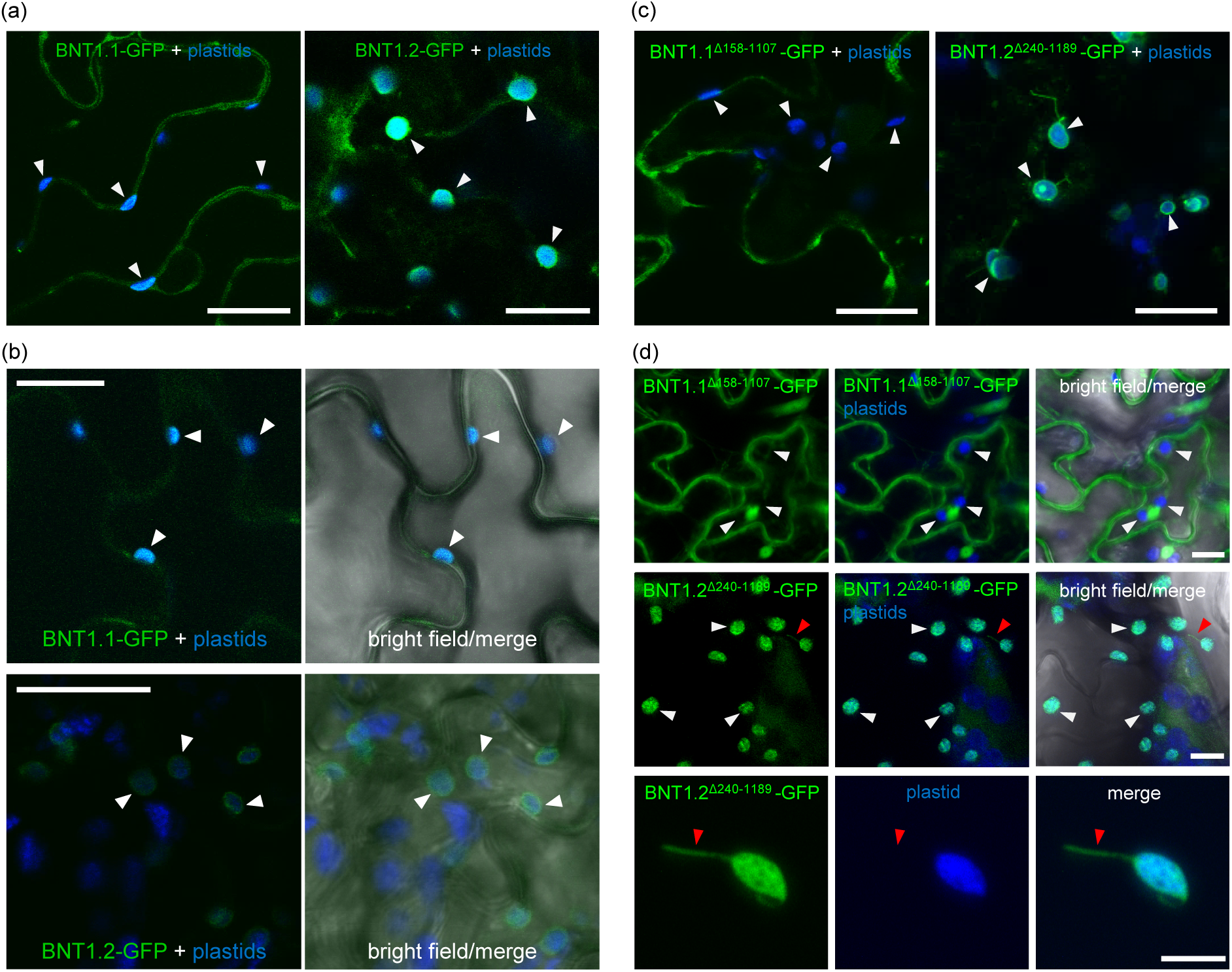
Differential subcellular localization of BNT1.1 and BNT1.2 isoforms. Confocal micrographs showing the subcellular localization of GFP-tagged full-length BNT1.1 and BNT1.2 isoforms (BNT1.1-GFP and BNT1.2-GFP) (a,b) and N-terminal domains (BNT1.1^Δ158-1107^-GFP and BNT1.2^Δ240-1189^-GFP). (a) Localization of BNT1.1-GFP and BNT1.2-GFP driven by the constitutive 35S promoter in transiently transformed *Nicotiana benthamiana* leaves. (b) Localization of BNT1.1-GFP and BNT1.2-GFP expressed under a Dex-inducible promoter in transgenic Arabidopsis 14-day-old seedlings. (c) Localization of N-terminal domains, BNT1.1^Δ158-1107^-GFP (TIR domain) and BNT1.2^Δ240-1189^-GFP (TM+cTP-like+TIR domains), driven by the 35S promoter in transiently transformed *N. benthamiana* leaves. (d) Z-series maximum intensity projections of adult transgenic *Arabidopsis* plants expressing N-terminal domains, BNT1.1^Δ 158-1107^-GFP and ^BNT1^.2^Δ240-1189^-GFP, under the 35S promoter. Bottom pane^ls: close^-up of BNT1.2^Δ240-1189^-GFP localization in a plastid envelope and stromule. Micrographs show GFP (green) and plastid autofluorescence (blue). White arrowheads: plastids, red arrowheads: stromules. In (a-c) Bar = 20 μm, and in (d) Bar = 10 μm.

These results indicate that the BNT1.1 isoform does not localize to plastids, confirming that the N-terminal signal is required for plastid envelope targeting. These findings also suggest that BNT1 isoforms function in distinct subcellular compartments, potentially contributing to different physiological processes.

### BNT1 is required for flg22-induced resistance against Pseudomonas

The expression analysis of total *BNT1*, *BNT1.1*, and *BNT1.2* isoforms (Figure 3 and Figure S4) suggests a potential role of BNT1 in PTI. Consequently, we evaluated BNT1 contribution to PAMPs-induced defense responses. To this end, we initially selected homozygous BNT1 T-DNA knockout mutant lines (Figure S5a-c). While previous studies have reported growth defects and flower sterility in *bnt1* mutant lines (Sarazin *et al*., 2015), we observed no significant differences in vegetative growth between WT Col-0 and mutant lines under our conditions (Figure S5d), suggesting that these phenotypes may depend on unknown environmental factors.

Next, we assessed the early apoplastic reactive oxygen species (ROS) burst and callose accumulation in response to the PAMP flg22 in WT Col-0 and *bnt1-7* mutant plants. As shown in Figure 5a and Figure S6a, the magnitude and kinetics of the flg22-induced ROS burst (0–30 min) were similar between *bnt1-7* and WT Col-0 plants. However, the number of callose deposits 24 h post-infiltration with 1 µM flg22 was significantly reduced in *bnt1-7* mutant compared to WT Col-0 plants (Figure 5b). Mock-treated leaves presented no differences between the two genotypes. These findings suggest a specific requirement for BNT1 in the late-stage of PTI responses (Nishimura *et al*., 2003; Wang *et al*., 2021). To test the susceptibility of the *bnt1-7* mutant to *Pst*, we employed two pathogen inoculation methods: leaf infiltration (OD_600_ = 0.005) and leaf spray (OD_600_ = 0.01) (Melotto *et al*., 2006; Zeng and He, 2010). Bacterial growth was not significantly different between *bnt1-7* and WT Col-0 plants with either method (Figure 5c and Figure S6b). Since *Pst* is highly virulent, we also tested the susceptibility of the *bnt1* mutant to *Pseudomonas syringae* pv. *tomato* DC3000 ΔhrcC (*Pst-*Δ*hrcC*). This strain is unable to inject effectors and, therefore useful to analyze PTI alone (Yuan and He, 1996; Hauck *et al*., 2003). Consistent with the results from virulent bacterial infections, no differences in *Pst-*Δ*hrcC* growth were observed between *bnt1-7* and WT Col-0 plants (Figure 5d).

**Figure 5.**
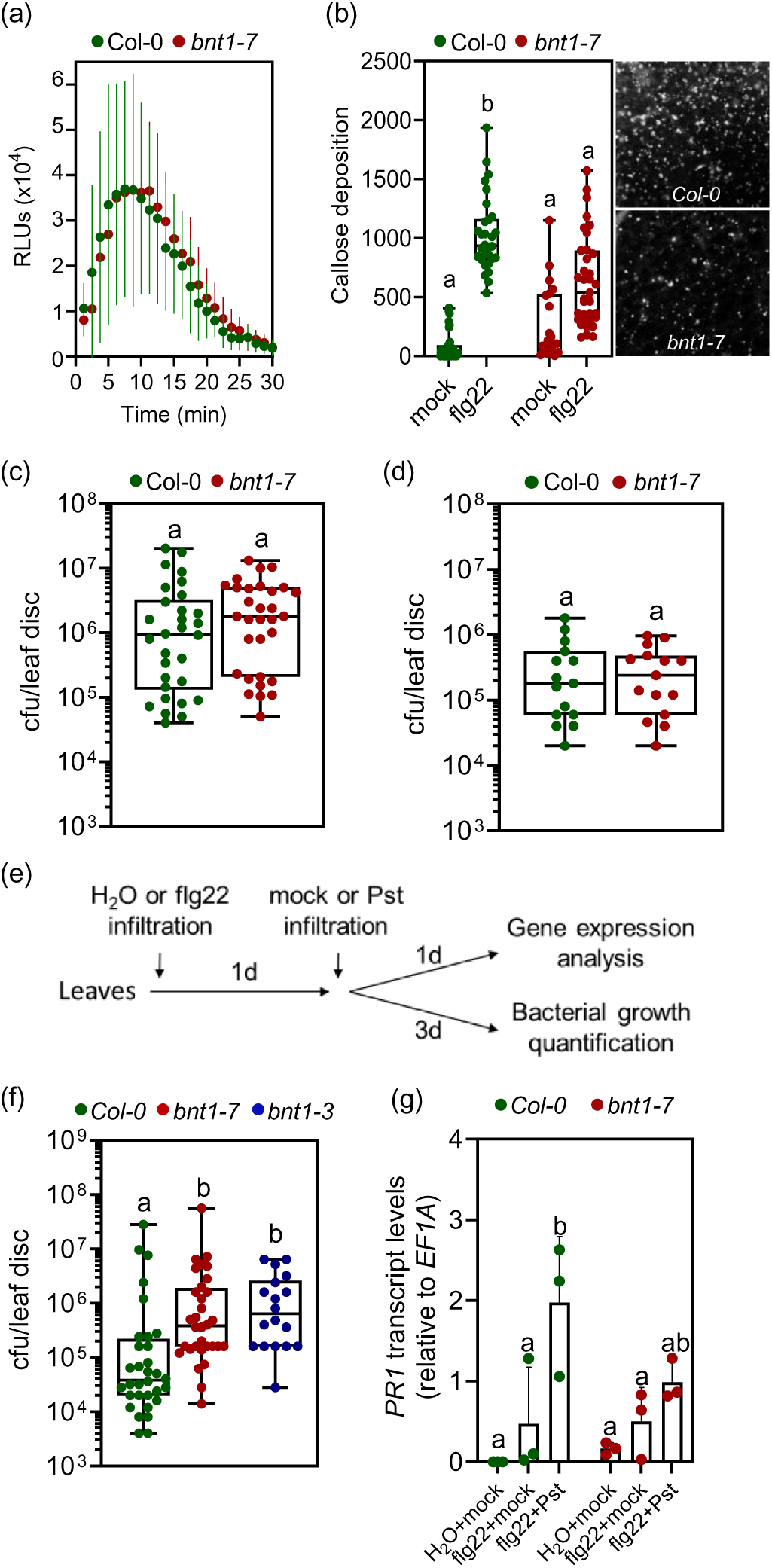
BNT1 is required for flg22-induced resistance against *Pst*. (a) Kinetics of flg22-induced apoplastic reactive oxygen species (ROS) production. Leaf discs from WT Col-0 and *bnt1-7* mutant plants were treated with mock or 100 nM flg22, and ROS was measured in relative light units (RLUs). The mean ± SD from one representative experiment is shown for each time point (leaf discs from at least 6 different plants; n = 24). (b) Number of callose depositions in WT Col-0 and *bnt1-7* leaves after 24 h post infiltration with mock (water) or 1 µM flg22. Right: representative micrographs showing aniline blue-stained callose deposits in flg22-infiltrated leaves. The number of deposits per image from two independent experiments (each one with at least 10 biological replicates; n = 20-35) is shown as scatter-dots in boxplots. (c) Growth of the virulent bacteria *Pst* on WT Col-0 and *bnt1-7* plants 3 days post-infection. *Pst* was inoculated by infiltration (OD_600_=0.005). The number of colony-forming units (cfu) per leaf disc is presented as scatter-dots in boxplots. Data from of four independent experiments is shown (each one with 6-8 biological replicates; n = 30). (d) Growth of the non-virulent bacteria *Pseudomonas syringae* pv. *tomato* DC3000 ΔhrcC (*Pst-*ΔhrcC) on WT Col-0 and *bnt1-7* plants 5 days post-infection. *Pst* was inoculated by infiltration (OD_600_=0.01). The number of cfu per leaf disc is presented as scatter-dots in the boxplots. The data from two independent experiments is shown (each one with 7-8 biological replicates; n=15). (e) Treatment scheme used in (f,g). (f) Growth of *Pst* on WT Col-0, *bnt1-7* and *bnt1-3* plants 3 days post infection. Leaves were inoculated with *Pst* (OD₆₀₀ = 0.005) 24 h after infiltration with 1 µM flg22, as shown in (e). The number of cfu per leaf disc is presented as scatter-dots in boxplots. Data from four independent experiments for Col-0 and *bnt1-7*, and two for *bnt1-3*, is presented (each one with 8 biological replicates; n=18-32) (g) Relative transcript levels of *PR1* in WT Col-0 and *bnt1-7* leaves, quantified by RT-qPCR. Leaves were treated with 1 µM flg22 or water, and after after 24 hpi infiltrated with Pst (OD₆₀₀ = 0.005) or mock (10 mM MgCl₂), as shown in (e). The mean ± SE from three independent experiments are shown (each data point with 3 leaves from 3 different plants pooled together for RNA extraction). (b-d,f,g) Different letters indicate statistically significant differences (p < 0.05, analysis of variance (ANOVA), Fisher’s LSD test).

We thus considered the possibility that BNT1 could be required for the later flg22-IR (Zipfel *et al*., 2004; Mishina and Zeier, 2007; De Kesel *et al*., 2021; Hönig *et al*., 2023). To study this, WT Col-0 and *bnt1-7* plants were pretreated with 1 µM flg22, and 24 h after, the same leaves were inoculated with *Pst* (OD_600_ = 0.005) (Figure 5e). Bacterial growth was assessed 3 days post-inoculation. Interestingly, the *bnt1-7* mutant showed increased susceptibility to *Pst* compared to WT Col-0 plants (Figure 5f). This phenotype was also observed in other allelic knockout line (*bnt1-3*; Figure 5f and Figure S4). Furthermore, in flg22-pretreated plants infected with *Pst* (Figure 5e), the defense marker gene *PR1* expression was lower in the *bnt1-7* mutant compared to WT Col-0 plants (flg22+*Pst*; Figure 5g). Conversely, no significant differences were observed in control or flg22-only treated plants (H_2_O+mock and flg22+mock; Figure 5g).

Together, these results indicate that BNT1 is important for callose accumulation and flg22-IR, positioning BNT1 as a downstream component of PTI likely contributing to the establishment of IR.

### Plastid-Targeted BNT1 mediates flg22-IR

The tissue-specific expression patterns and differential localization of BNT1.1 and BNT1.2 (Figures 3 and 4), together with the central role of plastids in plant defenses (Kachroo *et al*., 2021; Littlejohn *et al*., 2021; Yang *et al*., 2021) suggest that BNT1 isoforms may possess distinct immune functions. To explore this, we next studied their transcriptional behavior and involvement in flg22-IR.

We examined the changes in the transcript levels of *BNT1* isoforms associated to flg22-IR, using the experimental scheme described previously (Figure 5e). RT-qPCR analysis revealed a significant increase of the *BNT1.2* transcript abundance in response to *Pst* infection in flg22-pretreated plants (flg22+*Pst*) compared to control samples (H_2_O+mock and flg22+mock) (Figure 6a). Consistent with prior observations, *BNT1.1* exhibited substantially lower expression levels than *BNT1.2*, suggesting a predominant role for *BNT1.2* in flg22-IR. Nonetheless, a slight but significant increase in *BNT1.1* transcript levels was also detected under flg22+*Pst* treatment (Figure 6a).

**Figure 6.**
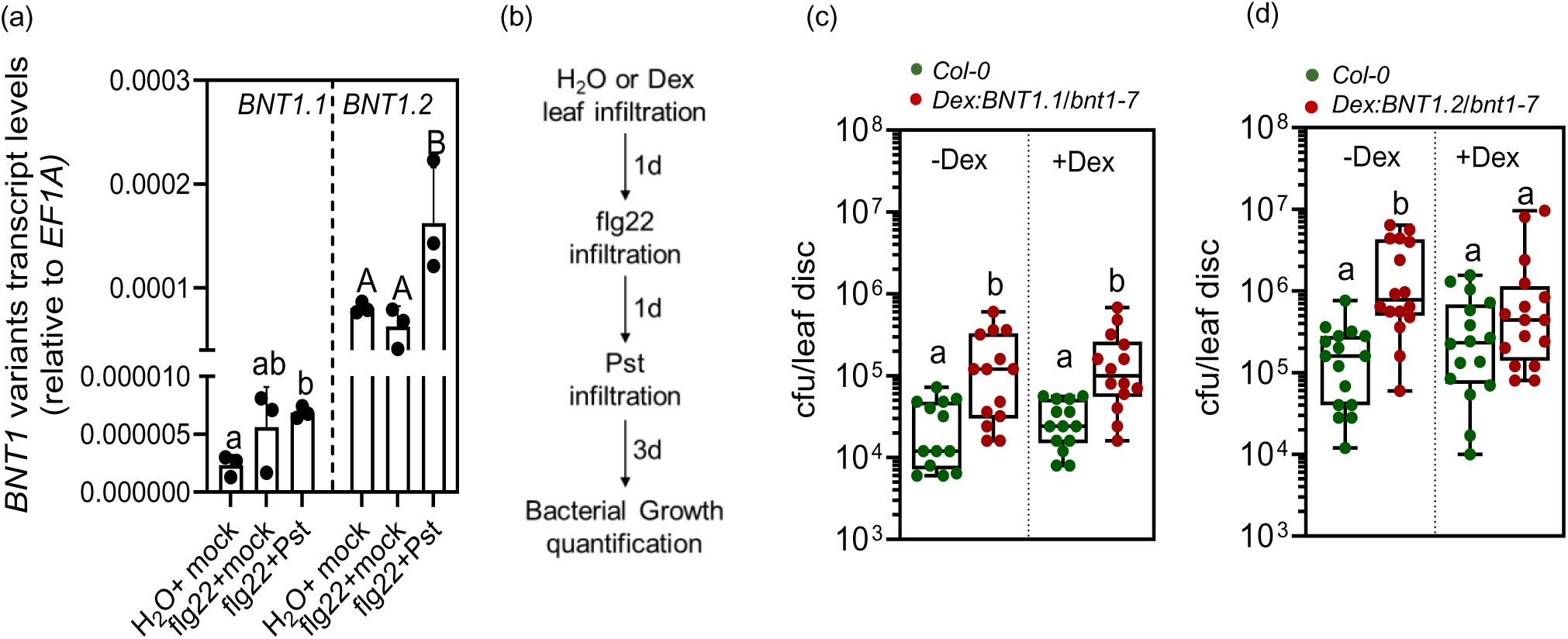
*BNT1.2* isoform rescues the flg22-induced resistance defective phenotype of *bnt1* mutant. (a) Relative transcript levels of *BNT1.1* and *BNT1.2* in WT Col-0 leaves, quantified by RT-qPCR using the specific primers indicated Figure 3a. Leaves were treated with 1 µM flg22 or water, and after 24 hpi infiltrated with *Pst* (OD₆₀₀ = 0.005) or mock (10 mM MgCl₂), as shown in Figure 5e. The mean ± SE from three independent experiments is shown (each data point with 3 leaves from 3 different plants pooled together for RNA extraction). Individual data points are presented as scatter-dots. Different letters (lowercase or uppercase) indicate statistically significant differences between treatments for each isoform (p < 0.05, ANOVA, Tukey test). (b) Treatment scheme used in (c,d). (c,d) To test the rescue of flg22-induced resistance defective phenotype, WT Col-0 or transgenic Dex:BNT1.1-GFP/*bnt1-7* (b) and Dex:BNT1.2:GFP/*bnt1-7* (c) plants were infiltrated with or without 30 µM dexamethasone (-Dex/+Dex) to induce *BNT1.1* or *BNT1.2* expression in *bnt1-7* mutant background. After 24 hours, the plants were treated with 1 µM flg22. *Pst* was inoculated into the same leaf one day later (OD=0.005) and bacterial growth was measured three days post-infection, as shown in (b). The cfu per leaf disc is presented as scatter-dots in the boxplots. Data from two independent experiments is shown (each one with at least seven biological replicates; n=14 (c) and n=16 (d)). Different letters indicate statistically significant differences between genotypes with or without Dex (p < 0.05, ANOVA, Fisher’s LSD test).

To further investigate the functional roles of *BNT1* isoforms, we performed flg22-IR assays against *Pst* using *bnt1* complementation lines expressing *BNT1.1-GFP* (*Dex:BNT1.1/bnt1-7*) or *BNT1.2-GFP* (*Dex:BNT1.2/bnt1-7*) under a Dex-inducible promoter (lines used in Figure 4b). WT Col-0 and the transgenic lines were treated with dexamethasone (+Dex) or mock (-Dex), infiltrated with 1 µM flg22, and 24 h later inoculated with *Pst* (OD_600_ = 0.005), as shown in Figure 6b. Bacterial growth was quantified 3 days post-infection. Both lines expressed the respective isoforms following Dex treatment (Figure S7). However, only the expression of *BNT1.2-GFP* successfully complemented the flg22-IR-deficient phenotype of the *bnt1-7* mutant, whereas *BNT1.1-GFP* did not (Figures 6c,d). To confirm the absence of complementation by *BNT1.1*, two additional independent *Dex:BNT1.1/bnt1-7* lines were examined, showing similar results (Figure S8).

These findings indicate that *BNT1.2* is sufficient to fulfill the role of *BNT1* in flg22-IR. Furthermore, they suggest that the functionality of this TNL in PTI requires plastid localization, highlighting a potential isoform-specific function for *BNT1*.

## DISCUSSION

Precise localization and trafficking of plant immune receptors are critical for their efficiency (Padmanabhan and Dinesh-Kumar, 2010; Qi and Innes, 2013; Bernoux *et al*., 2023; Shepherd *et al*., 2023). Here, we identified an alternative TSS variant of the TNL-class NLR immune sensor BNT1 at the plastid envelope, where it contributes to PTI. This finding is noteworthy in light of the fact that plastid-targeted NLRs (and other immune receptors) have remained largely unexplored, despite the central role of plastid-dependent defense responses (Kachroo *et al*., 2021; Littlejohn *et al*., 2021; Yang *et al*., 2021; Breen *et al*., 2022). Our results highlight the importance of the subcellular NLRs localization and reinforce the central role of plastids in plant immunity.

### Plastid-envelope targeting of BNT1

Our biochemical and microscopy data indicate that BNT1’s plastid envelope localization relies on the signal-anchored (SA) protein targeting mechanism (Figure 1 and 2). SA proteins possess an N-terminal non-cleavable transmembrane domain (TM) that anchors them to the membranes of the ER, plastids, or mitochondria, depending on the TM hydrophobicity (Kim and Hwang, 2013). The BNT1.2 protein features a bipartite N-terminal signal (TM+cTP-like), representing a variation of the SA-targeting protein motif, that enables ER-predicted SA proteins to target plastid envelopes (Banday *et al*., 2022). However, the cTP-like domain in BNT1.2 is not strictly required for plastid targeting, as shown for other defense factors (Cecchini *et al*., 2015b). Instead, it appears to induce stromule formation and plastid clustering around nuclei when ectopically expressed in *N. benthamiana*, both of which are features of defense induction in different pathosystems (Liu *et al*., 2024). Thus, while a complete signal is unnecessary for BNT1 localization, it could influence plastid behavior in response to pathogen attacks. Supporting this idea, the cTP-like domain in the bipartite signal of the defense factor AZI1, was recently shown to associate with microtubules (Cecchini *et al*., 2021), which play a central role in stromule extension and plastid clustering (Erickson *et al*., 2017; Erickson and Schattat, 2018; Kumar *et al*., 2018; Hanson and Conklin, 2020; Jung *et al*., 2024). Future work will address this possibility.

A limited number of NLRs have been identified with N-terminal TM signal anchors determining their localization to specific membranes (Takemoto *et al*., 2012). However, given the ongoing challenges in recognizing TMs that target plastid and mitochondrial envelopes (Lee *et al*., 2011; Kim and Hwang, 2013; Cecchini *et al*., 2021; Banday *et al*., 2022), the number of SA NLRs in endosymbiont organelles is likely underestimated. Supporting this idea, *in silico* screenings have predicted several well-known and uncharacterized plastid– and mitochondria-targeted SA proteins, including NLRs and NLR-related proteins (Lee *et al*., 2011; Banday *et al*., 2022). Among these, experimental evidence indicates that *Arabidopsis* RPW8 family members HR2 and HR3 localize at plastids (Lee *et al*., 2011; Berkey *et al*., 2017). Additionally, RPW8.1 was previously detected in chloroplasts (Wang *et al*., 2007; Ma *et al*., 2014). Since the RPW8-like CC domain is a characteristic feature of helper CC_R_-NLRs, such as *N. benthamiana* NRG1, which—together with the TNL BNT1—are the only two NLRs currently known to localize to plastid envelopes (Ibrahim *et al*., 2024), further localization studies will probably uncover additional signal-anchored CNLs and TNLs in endosymbiont organelles. Moreover, it is also possible that ER-anchored NLRs could relocate under specific defense conditions, as observed for other receptors and key defense factors (Kim *et al*., 2014; Medina-Puche *et al*., 2020; Duggan *et al*., 2021).

### TSS-driven specific locations of BNT1 isoforms

Fine-tuning gene expression is crucial for plant environmental adaptability (Fick *et al*., 2022; Zhong *et al*., 2023). One such mechanism, the use of alternative TSSs, has been shown to optimize gene function by modifying 5′ UTR sequences. They influence mRNA stability, translational efficiency, and/or protein localization (de Klerk and ‘t Hoen, 2015; Zhang *et al*., 2022). Notably, we found that through alternative TSS usage, the *BNT1* gene produces two distinct isoforms: BNT1.2, localized to the plastid envelope, and BNT1.1, targeted to the cytoplasm (Figures 3 and 4). Since these isoforms are differentially regulated, with *BNT1.2* expression significantly upregulated and *BNT1.1* downregulated following *Pst* infection, BNT1 may function in distinct subcellular compartments, fulfilling specific roles in plant immunity. The reprogramming of TSS usage likely enriches BNT1 at the plastid envelope, suggesting that this location is required during the interaction between *Arabidopsis* and *Pst*. This is further supported by the observation that only BNT1.2, not BNT1.1, is necessary for the flg22-IR against *Pst* (Figure 6). This is consistent with recent findings demonstrating extensive TSS reprogramming during PTI (Thieffry *et al*., 2022) and the requirement of a plastid-targeted, alternative TSS-generated glycerate 3-kinase for ETI (Gao *et al*., 2020).

Additionally, *BNT1.2* was predominantly expressed in epidermal cells, whereas *BNT1.1* appeared to show higher expression in vascular and mesophyll cells, suggesting tissue-specific expression of *BNT1* driven by alternative TSSs. Consistently, BNT1 peptides have been identified in plastids of epidermal and vascular parenchyma cells (Beltrán *et al*., 2018). This suggests that the role of BNT1.2 in flg22-IR is specific to plastids in epidermal cells. The leaf epidermis is a key site for pathogen recognition and the initiation of both local and systemic defense responses, including IR programs (Zheng *et al*., 2015; Vlot, 2021; Irieda and Takano, 2021; Jiang *et al*., 2021; Irieda, 2022). Moreover, epidermal plastids, referred to as “sensory plastids”, are proposed to be specialized organelles for detecting abiotic and biotic stresses and initiating retrograde signaling to the nucleus (Mackenzie and Mullineaux, 2022; Sierra *et al*., 2023). Thus, NLRs like BNT1 and other resistance proteins localized in sensory plastids may contribute to the unique characteristics of these epidermal organelles by facilitating pathogen perception and/or signaling defense responses. Supporting this idea, members of the RPW8/HR family are differentially expressed between mesophyll and epidermal tissues and localize to plastid envelopes, the plasma membrane, and/or the extrahaustorial membrane (Wang *et al*., 2007; Wang *et al*., 2009; Ma *et al*., 2014; Berkey *et al*., 2017). Future research into tissue-specific expression patterns of NLRs will provide valuable insights into their distinct roles in plant immune responses.

### The role of BNT1.2 in PTI responses

NLRs are typically associated with ETI, where they detect pathogen effectors (Jones *et al*., 2016). However, recent studies have revealed that PTI and ETI are interconnected, sharing components and reinforcing each other (Jones *et al*., 2024; Ngou *et al*., 2021; Bernoux *et al*., 2022). Here we showed that BNT1 is specifically required for certain PTI-associated responses, such as callose deposition, while it is dispensable for early apoplastic ROS production (Figure 5). Additionally, BNT1 is involved in flg22-IR against *Pst* and thus, potentially in PAMP-induced defense priming (Zipfel *et al*., 2004; Mishina and Zeier, 2007; Martinez-Medina *et al*., 2016). Supporting this, the *bnt1* mutant fails to show a boosted *PR1* expression in response to *Pst* after flg22 pre-treatment but exhibits WT-like *PR1* levels under control conditions, a hallmark of the immune memory or priming (De Kesel *et al*., 2021). Interestingly, only the plastid-localized BNT1.2 isoform is essential for flg22-IR (Figure 6), suggesting that the right compartment localization is essential for BNT1 participation in PTI responses. Additionally, *BNT1.1* appears to be more abundant in the vasculature and mesophyll, while *BNT1.2* in epidermal cells. This differential spatial distribution, if it correlates with protein levels, could also provide a rationale for the distinct functions of the two isoforms. It is plausible that BNT1.1 plays a defense role against pathogens that target vascular tissues. Alternatively, BNT1.1 retains all key residues necessary for its activity (Krasileva *et al*., 2010; Toshchakov and Neuwald, 2020) but lacks the first conserved β-sheet (βA) of the TIR domain (Maruta *et al*., 2022), which may influence its functionality or interactions (Bernoux *et al*., 2011; Ma *et al*., 2020; Martin *et al*., 2020). Future experiments will elucidate the distinct functions BNT1.1 and BNT1.2.

Thus, what is BNT1’s role in plastid envelopes? One idea is that it senses plastid changes associated with PTI. Supporting this, it was demonstrated that PAMPs perceived at the plasma membrane are known to induce a specific Ca²⁺ increase in the stroma and chloroplast ROS production or redox state changes (Nomura *et al*., 2012; Zabala *et al*., 2015; Arnaud *et al*., 2022). Moreover, these fluctuations are needed for PAMP-triggered callose deposition (Nomura *et al*., 2012). Therefore, an NLR located within the plastids appears to be optimally positioned to detect the organelle alterations. We propose that BNT1 TNL guards unknown component(s)/metabolite(s) in the sensory plastids. Alternatively, BNT1 activation could modulate plastid-associated PTI responses. Given the prevalence of pathogen effectors targeting plastids to promote virulence, BNT1 may also be sensing effector-induced organelle disruptions.

Furthermore, BNT1 activation might recruit immune complexes, like the EDS1-lipase-like proteins and CC_R_-NLR helpers (Gong *et al*., 2023; Shepherd *et al*., 2023), to signal downstream defenses. In agreement, PTI-associated responses depend on the EDS1-PAD4/SAG101 and ADR1s/NRG1s pathways (Tian *et al*., 2021; Pruitt *et al*., 2021). However, in *Arabidopsis*, these factors are localized to the plasma membrane, ER, or cytoplasm (Wu *et al*., 2019; Saile *et al*., 2021; Feehan *et al*., 2023). Therefore, one possibility is that BNT1 activation induces the relocalization of the complex components to the plastid envelope. Alternatively, the plastid-localized RPW8/HR family members (Lee *et al*., 2011; Berkey *et al*., 2017) might function in association with BNT1, as shown for HR4 and the RPP7 NLR (Li *et al*., 2020). If either scenario holds true and considering that plastids are intracellular calcium stores (Stael *et al*., 2012), it is conceivable that during PTI, CC_R_-NLRs (or RPW8/HRs) could form pores across the organelle double membranes, as has been suggested for *N. benthamiana* NRG1 (Ibrahim *et al*., 2024). Consequently, BNT1 activation may increase cytoplasmic calcium, potentially linking PTI-mediated plastid fluctuations to retrograde defense signaling.

Altogether, our findings highlight a potentially central role for plastid-targeted NLRs in plant immunity. We speculate that NLRs in plastids (and mitochondria) might act as nodes integrating PTI and ETI signaling. However, further exploration is needed to identify additional NLRs in endosymbiont organelles, elucidate their functions in distinct tissues, and uncover their inter-compartmental signaling mechanisms. Future research will be crucial to understand the role of plastids in hosting immune receptors and mediating plant defense.

## EXPERIMENTAL PROCEDURES

### Plants and vectors

All *Arabidopsis thaliana* plants used were Columbia-0 (Col-0) ecotypes. Mutant lines *bnt1-3* (SALK_082608C) and *bnt1-7* (GK-392D02) were obtained from the *Arabidopsis* Biological Resource Center (Ohio State University, Columbus, OH, USA). Transgenic plants expressing the GFP-tagged BNT1.1 and BNT1.2, or their N-terminal regions, were generated by the floral dip method (Clough and Bent, 1998). Transformations were done with *Agrobacterium tumefaciens* strain GV3101 harboring the respective constructs. Transformed plants were selected with Basta (10 µg ml^-1^) or kanamycin (50 µg ml^-1^), and F2/F3 generation plants were used for experiments. *Arabidopsis* were grown under a 12-hour light/12-hour dark photoperiod at 20-22°C, 100-120 µmol sec^-1^ m^-2^ light intensity, and 50-70% relative humidity. Adult plants used in the experiments were 4 weeks old. Seedlings were grown on ½ Murashige-Skoog medium containing 0.8% agar and 1% sucrose for 14 days. *Nicotiana benthamiana* plants utilized for *Agrobacterium tumefaciens*-mediated transient transformation were grown at 22-24°C under a 16-hour light/8-hour dark photoperiod for 4-6 weeks.

The full-length *BNT1.1* and *BNT1.2* coding sequences, their N-terminal regions, or the bipartite signal variants were amplified and cloned into the TOPO pENTR® vector. These constructs were then transferred to the plant expression vectors pBAV150 ((Dex)-inducible promoter, C-terminal GFP-tagged; (Vinatzer *et al*., 2006)) and/or pGWB405 (35S constitutive promoter, C-terminal GFP-tagged; (Nakagawa *et al*., 2007)) using GATEWAY® technology. The chloroplast outer envelope protein marker OEP7 (OEP7:RFP) and the ER marker BiP (BiP:RFP) fused to RFP were previously described (Cecchini *et al*., 2015b). All primers, vectors, and constructs used in this study are listed in Table S1.

### Subcellular localization

For localization studies in *N. benthamiana*, leaves were infiltrated with *A. tumefaciens* carrying the different vectors. For constructs under the control of dexamethasone (Dex)-inducible promoters, *A. tumefaciens* was inoculated into leaves and, after 24 h, infiltrated with 30 μm dexamethasone (Dex) (D4902; Sigma-Aldrich). Fractionation and confocal microscopy studies were conducted 21 h after Dex treatment. *N. benthamiana* leaves were transformed with vectors using the constitutive 35S promoter, and confocal imaging was carried out 36 hours post-agroinfiltration. Transgenic *Arabidopsis* seedlings were immersed in 30 μM Dex solution for 16 hours before imaging.

As described previously, leaf discs from *N. benthamiana* and *Arabidopsis* were prepared for microscopic analysis (Cecchini *et al*., 2022). Imaging was carried out using Olympus FV1000 or FV1200 laser scanning spectral confocal microscopes (Olympus, Latin America) to detect GFP (excitation: 488 nm; emission: 505–530 nm), RFP (excitation: 561 nm; emission: 570–620 nm), and plastid autofluorescence (excitation: 633 nm; emission: 650–750 nm). Images were captured in sequential acquisition mode using LD C-Apochromat 40×/1.1 W Korr or PLAPON 60×/1.42 AN objective (Olympus, Japan). Time-lapse series and Z-stack scans were acquired using lower scanning resolution and maximum speed settings. Images were processed with Fiji software (https://fiji.sc/) and Adobe Photoshop®. All microscopy analyses were conducted at the Center for Microscopy and Nanoscopy of Córdoba (CEMINCO; https://ceminco.conicet.unc.edu.ar/).

### Fractionations, membrane association and Western blot analysis

Microsomal and plastid fractions were isolated as previously (Cecchini *et al*., 2015b; Cecchini *et al*., 2021), using 0.5 g of *N. benthamiana* leaf tissue expressing the different constructs. Membrane association assays were conducted with microsomal extracts. Briefly, the extracts were treated with 1.5 M NaCl, 2 M urea, 1% Triton X-100, or 1% NP-40 + 0.5% deoxycholate, followed by gentle shaking at room temperature for 20 minutes. The samples were then ultracentrifuged to separate soluble supernatants from membrane pellet fractions. Soluble fractions were precipitated with trichloroacetic acid before immunoblotting.

For Western blotting, total proteins or fractionated samples were separated by SDS-PAGE. Protein concentrations were determined using the Bradford assay (Bradford, 1976). Primary anti-GFP antibodies used for detection included Covance MMS-118P (1:5000) and Roche #10744900 (1:3000). Secondary antibodies used were either horseradish peroxidase-conjugated anti-mouse antibodies (Thermo Scientific, Rockford, IL, USA, 1:1000) or infrared fluorescent anti-mouse antibodies (LI-COR 926–32 211 and 925–32 210, 1:25000). Band detection was performed using SuperSignal Stable Peroxidase (Thermo Scientific) or an Odyssey Infrared scanner (LI-COR Biosciences), respectively.

### Tissues Isolation and Enrichment

Mesophyll, epidermal, and vascular bundle tissues were isolated following the protocols of (Endo *et al*., 2016) with minor modifications. Between 5 and 10 mature *Arabidopsis* leaves were placed between layers of labeling tape, and the abaxial epidermis was carefully peeled away. The epidermal peels and remaining leaf tissues were transferred to separate 60 × 15 mm Petri dishes containing 3 mL of enzyme digestion buffer (0.4 M mannitol, 10 mM CaCl_2_, 20 mM KCl, 0.1% [wt/vol] BSA, 20 mM MES, pH 5.7, 1% [wt/vol] cellulase, 0.25% [wt/vol] macerozyme). After 15 minutes of incubation at room temperature, vascular bundles were isolated from the remaining tissues using fine-tipped forceps, washed in washing buffer (154 mM NaCl, 125 mM CaCl_2_, 5 mM KCl, 2 mM MES, pH 5.7), and flash-frozen in liquid nitrogen for RNA extraction. The epidermal peels were washed and incubated in digestion buffer for an additional 10 minutes to release epidermal cells from the tape. The resulting cell suspension was centrifuged at 100 × *g* for 5 minutes at 4°C, and the pelleted cells were resuspended in washing buffer. The remaining tissues underwent further digestion for 40 minutes to release mesophyll cells. These cells were collected, centrifuged at 100 × *g* for 5 minutes at 4°C, and resuspended in washing buffer. Both mesophyll and epidermal cells were then centrifuged and washed under the same conditions, with the final pellets flash-frozen for RNA extraction.

RNA samples were analyzed to confirm tissue-specific enrichment. The expression levels of tissue-specific marker genes (Table S1) were evaluated to validate the isolation procedure (Figure S3b; (Endo *et al*., 2016)).

### Apoplastic ROS measurement

flg22-induced apoplastic ROS accumulation was measured as described (Fabro *et al*., 2016), with minor modifications. Leaf discs (0.38 cm²) were excised from adult plants using a cork borer and transferred into 96-well plates containing 150 µL of water per well using a fine paintbrush. The plates were kept in darkness for 16 hours. Three hours before luminescence measurements, the discs were washed with water. A reaction mixture containing 100 nM flg22, 34 μg/mL Type IV horseradish peroxidase, and 20 μg/mL luminol were added to each well, and luminescence was immediately quantified using a Synergy HT plate reader (Biotek). Measurements were recorded every 73 seconds for 30 minutes, using an integration time of 0.3 seconds per well and a gain of 180. The flg22 peptide (QRLSTGSRINSAKDDAAGLQIA) was obtained from GenScript (RP19986), dissolved in Milli-Q water to prepare a 1 mM stock solution, and stored in 20 µL aliquots at –20°C until use.

### Callose quantitation

Callose was quantified using aniline blue staining (Kim and Mackey, 2008). Leaves were collected 24 hours after treatment with 1 µM flg22 or water infiltration, and chlorophyll was cleared with 95% ethanol. Measurements were taken from at least 10 leaves collected from different plants for each genotype and treatment. Images of callose deposits were captured using an Axioplan epifluorescence microscope (Zeiss, Germany) equipped with a DAPI filter set, and deposits were quantified using the Fiji-TWS method (Zhou *et al*., 2012). Results are expressed as the number of callose deposits per image.

### Pathogen infection and flg22-induced resistance

To evaluate bacterial growth in wild-type (WT Col-0), mutant, or transgenic *Arabidopsis* lines, leaves from 4-week-old plants were infected either by syringe infiltration (OD_600_ = 0.005) or by spraying (OD_600_ = 0.01) with virulent *Pseudomonas syringae* pv. *tomato* DC3000 (*Pst*; (Ritter and Dangl, 1996)) or the mutant strain *Pseudomonas syringae* pv. *tomato* DC3000 ΔhrcC (*Pst-*ΔhrcC) (Yuan and He, 1996; Hauck *et al*., 2003). Bacterial growth was quantified by dilution plating, counting CFUs on King’s B agar plates after 3 days at 28°C.

For local flg22-induced resistance (flg22-IR) assays, *Arabidopsis* leaves were infiltrated with 1 µM flg22. After 24 hours, the same leaves were infiltrated with Pst (OD_600_ = 0.005). Bacterial growth was measured 3 days later. For gene expression analysis, leaves were harvested and frozen 1-day post-infection for RNA extraction. To analyze flg22-IR rescue of the *bnt1-7* mutant by BNT1.1 or BNT1.2, transgenic complementation lines Dex:BNT1.1-GFP (*bnt1-7*) or Dex:BNT1.2-GFP (*bnt1-7*) were infiltrated with water or 30 µM dexamethasone 24 hours before 1 µM flg22 treatment. One day after flg22 infiltration, leaves were inoculated with Pst (OD_600_ = 0.005), and bacterial growth was quantified 3 days after.

### Transcripts level analysis

Total RNA was extracted using Trizol and reverse-transcribed using MMLV enzyme according to the manufacturer’s instructions (M-MuLV reverse transcriptase, NEB #M0253S). Quantitative real-time RT-PCR (RT-qPCR) was performed as previously described (Miranda de la Torre *et al*., 2023), using a CFX96 Real-Time PCR Detection System (Bio-Rad) or Real-Time PCR AriaMx (Agilent). Raw data were baseline-corrected, and linearity was assessed using LinRegPCR 2021.1 (Ruijter *et al*., 2009). The *Elongation Factor 1 alpha* gene (*EF1*α, *At5g60390*) and *YELLOW-LEAF-SPECIFIC GENE 8* (*YLS8*, *At5g08290*) were used as reference genes (He *et al*., 2017). *BNT1.1* and *BNT1.2* expression analysis by semi-quantitative RT-PCR (RT-sqPCR) was tested by a 35 cycle PCR. Sequences of all the primers used in this study are shown in Table S1.

*BNT1* levels under biotic stress conditions and different tissues were obtained from Genevestigator® (Perturbations and Anatomy tools; https://genevestigator.com/). Only perturbations with significant fold-changes (nominal *t*-tests, p < 0.002, fold-change > 2) were considered.

### Statistical Analysis

Analyses in this study were performed using GraphPad Prism 10 and InfoStat v2018 (www.infostat.com.ar; Grupo InfoStat). Grub’s test identified and excluded outliers (α = 0.05). Square root-transformed data were applied for bacterial growth curves in ANOVA tests. Analysis of variance (ANOVA), followed by post hoc tests, was conducted to determine significant differences, as specified in the figure legends.

## DATA AVAILABILITY STATEMENT

All relevant data can be found within the manuscript and its supporting materials.

## ACKNOWLEDGMENTS

This research was supported by grants from ANPCyT (PICT-2020-01906 to MPM, PICT-2017-0589 and PICT-2020-0483 to NMC, PICT-2018-4588 and PICT-2021-00805 to MEA), SECyT-UNC (to MEA), CONICET (PIP-2021-2023 to NMC), NSF (IOS1456904 to Jean T. Greenberg), ANID (FONDECYT Regular 1210320 to FBH, FONDECYT Postdoc 3240308 to MPM), PIA/BASAL AFB240003 and Programa Iniciativa Científica Milenio— (IBio) ICN17_022 to FBH. MPM received support from an ICGEB SMART Fellowship, a CONICET postdoctoral fellowship, and an ANID fellowship. NMC and MEA are Career Investigators of CONICET.

We especially thank Dr. Jean T. Greenberg at The University of Chicago, where this story began, for her insightful discussions and continued support over the years. We also sincerely thank Dr. Ezequiel Petrillo for his assistance with TSS-related experiments and for sharing unpublished data, as well as Dr. Georgina Fabro and Dr. Suruchi Roychoudhry for their critical and thorough review of the manuscript. Our gratitude extends to the members of the NMC, MEA, FBH and Georgina Fabro labs for their valuable discussions. We further thank to Dr. María Florencia Nota (professional of CIQUIBIC-CONICET) for technical assistance and to Dr. Carlos Mas and Dr. Cecilia Sampedro (professionals at the Centro de Micro y Nanoscopía de Córdoba [CEMINCO]) for their help with confocal microscopy.

## AUTHOR CONTRIBUTIONS

MPM and NMC conceived and designed the experiments. MPM, NMC, AH, APC, JRP and FB performed the experiments. MPM, NMC, FBH, AH, and MEA analyzed and interpreted data. MPM and NMC wrote the paper. All the authors read, reviewed, and edited the final manuscript.

## SUPPORTING INFORMATION

Additional Supporting Information may be found in the online version of this article.

**Figure S1**. Localization of BNT1-GFP in leaves of transgenic *Arabidopsis thaliana* plants.

**Figure S2**. Alternative transcription start sites (TSSs) generating *BNT1.1* and *BNT1.2* transcripts.

**Figure S3**. Alternative *BNT1* transcripts under basal conditions.

**Figure S4**. Expression data for BNT1 (AT5G11250).

**Figure S5**. Characterization of T-DNA insertion mutant lines.

**Figure S6**. Total apoplastic reactive oxygen species (ROS) and growth of sprayed *Pst* in WT (*Col-0*) and *bnt1-7* mutant plants.

**Figure S7**. Characterization of *Arabidopsis* transgenics lines Dex:BNT1.1-GFP/*bnt1-7* (*line #3*) and Dex:BNT1.2:GFP/*bnt1-7* (*line #1*).

**Figure S8**. Characterization of *Arabidopsis* transgenics lines Dex:BNT1.1-GFP/*bnt1-7*

**Table S1.** Vectors, constructs and primers list used in this study

**Movie S1.** Localization of BNT1-GFP in *N. benthamiana* epidermal cell plastids and stromules surrounding a nucleus.

**Movie S2.** Localization of BNT1-GFP in plastids of *Arabidopsis* epidermal cells.

