## Supplementary material for "The *Arabidopsis* TNL immune receptor BNT1 localizes to the plastid envelope and mediates flg22-induced resistance against *Pseudomonas*": Figure S1

(a)

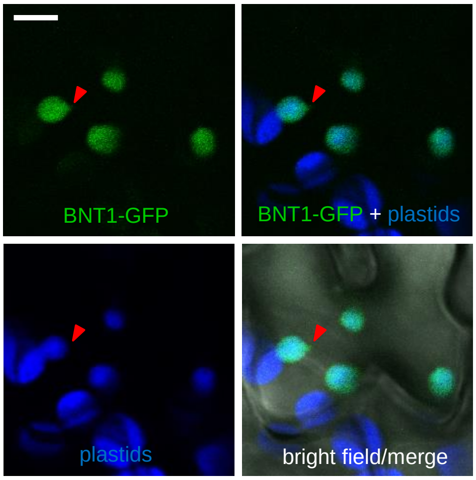


**Figure S1.** Confocal micrographs showing BNT1-GFP in leaves of transgenic *Arabidopsis thaliana* plant. Red arrowheads: stromule. Asterisks indicate spongy mesophyll cells plastids. Micrographs show GFP (green), RFP (red) and plastid autofluorescence (blue). Bar = 5 μm.
