## Supplementary material for "The *Arabidopsis* TNL immune receptor BNT1 localizes to the plastid envelope and mediates flg22-induced resistance against *Pseudomonas*": Figure S2

(a)

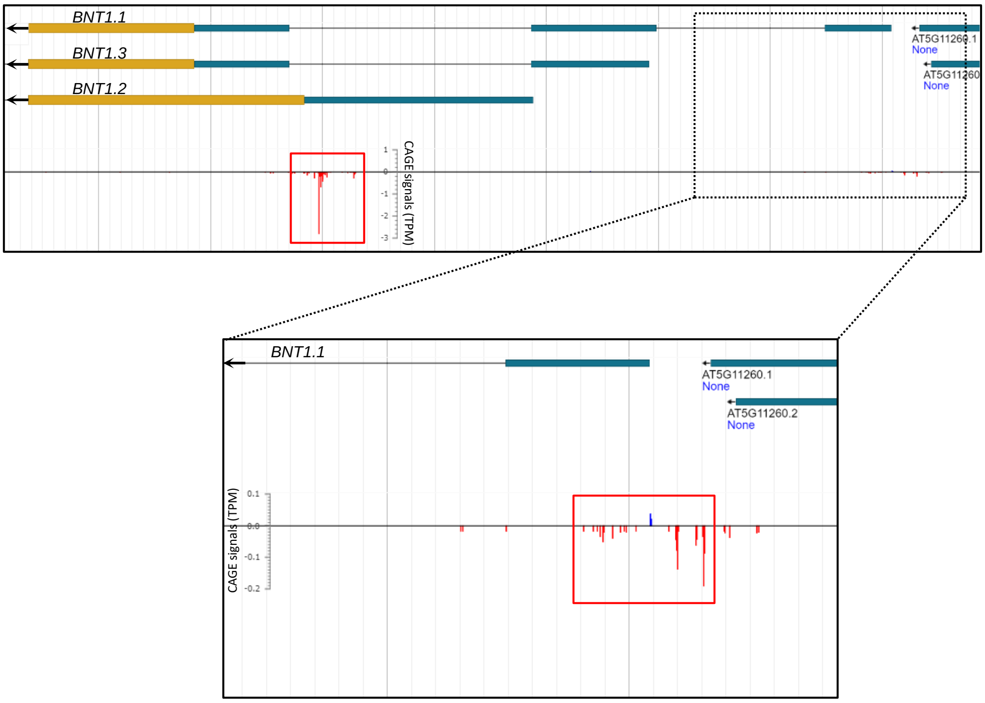


(b)

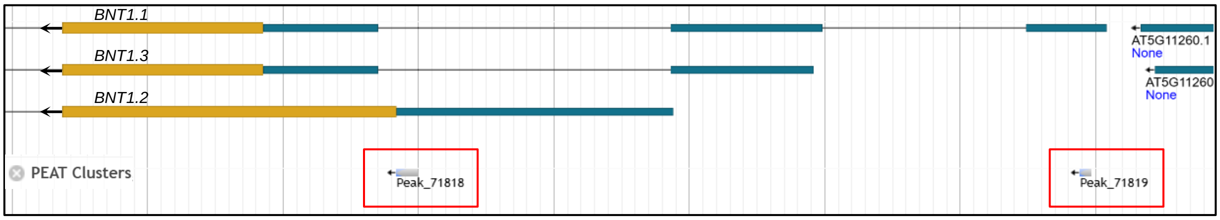


**Figure S2.** Alternative transcription start sites (TSSs) generating *BNT1.1* and *BNT1.2* transcripts.
(a,b) Diagrams showing the three predicted alternative transcripts (*BNT1.1*, *BNT1.2*, and *BNT1.3*) of the *BNT1* gene model in Araport11. Yellow blocks: coding exons, blue blocks: 5' UTR exons, black lines: introns.
(a) JBrowse track displaying CAGE (Cap Analysis of Gene Expression) normalized signals (TPM, transcript per million) from three biological replicates of 14-day-old WT Col-0 seedlings under basal conditions (wt_R123, 0 min; Thieffry et al., 2020). Red boxes indicate clusters of CAGE tags for *BNT1.1* and *BNT1.2*. Plus strand: blue, minus strand: red. Bottom: Close up from genome region (dashed box) showing BNT1.1 CAGE tags cluster. (https://jbrowse.arabidopsis.org/index.html?data=Araport11&loc=Chr5%3A3591029..3593896&tracks=TAIR10_genome%2CA11-PC%2Cwt_R123%20(0%20min)&highlight=).
(b) JBrowse track showing PEAT (paired-end analysis of transcription start sites) clusters from 10-day-old WT Col-0 seedling roots under basal conditions (Morton et al., 2014). Red boxes indicate *BNT1.1* and *BNT1.2* PEAT peaks (each PEAT peaks show a minimum of 10 reads). (https://jbrowse.arabidopsis.org/index.html?data=Araport11&loc=Chr5%3A3591055..3593660&tracks=TAIR10_genome%2CA11-GL%2CA11-PC%2CAll_PEAT&highlight=).
Schemes were adapted from Arabidopsis JBrowse 1.16.6; JBrowse 2: a modular genome browser with views of synteny and structural variation. Genome Biology (2023). https://doi.org/10.1186/s13059-023-02914-z).
