## Supplementary material for "The *Arabidopsis* TNL immune receptor BNT1 localizes to the plastid envelope and mediates flg22-induced resistance against *Pseudomonas*": Figure S3

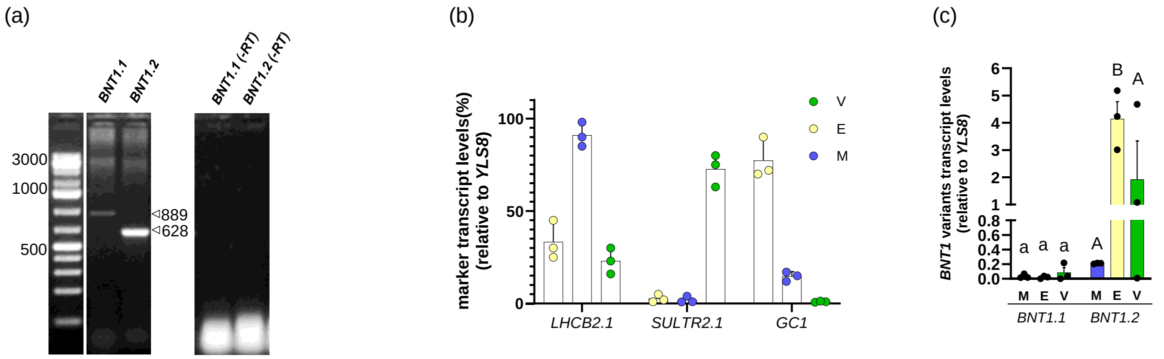


**Figure S3.** Alternative *BNT1* transcripts under basal conditions.
(a) RT-PCR analysis showing negative controls (-RT) from the experiments in Figure 3c. Arrowheads indicate expected band sizes (*BNT1.1:* 889 bp, *BNT1.2*: 628 bp).
(b) Transcript levels of marker genes for isolated mesophyll cells (*Lhcb2.1*), vascular bundle (*Sultr2;1*), and epidermal cells (*GC1*), showing tissue-specific enrichment from *Arabidopsis* leaves used in Figure 3d. M: mesophyll; E: epidermis; V: vascular bundle. The mean ± SE from three independent experiments is shown (each data point with 6-8 leaves from at least 3 different plants pooled together for RNA extraction).
(c) Basal transcript levels of *BNT1.1* and *BNT1.2* in different leaf tissues (M: mesophyll; E: epidermis; V: vascular bundle) under basal conditions. The mean ± SE from three independent experiments is shown (each data point with 6-8 leaves from at least 3 different plants pooled together for RNA extraction). Different letters (lowercase or uppercase) indicate statistically significant differences between tissues for each isoform ($p < 0.05$, ANOVA, Fisher´s test).
