## Supplementary material for "The *Arabidopsis* TNL immune receptor BNT1 localizes to the plastid envelope and mediates flg22-induced resistance against *Pseudomonas*": Figure S4

(a)

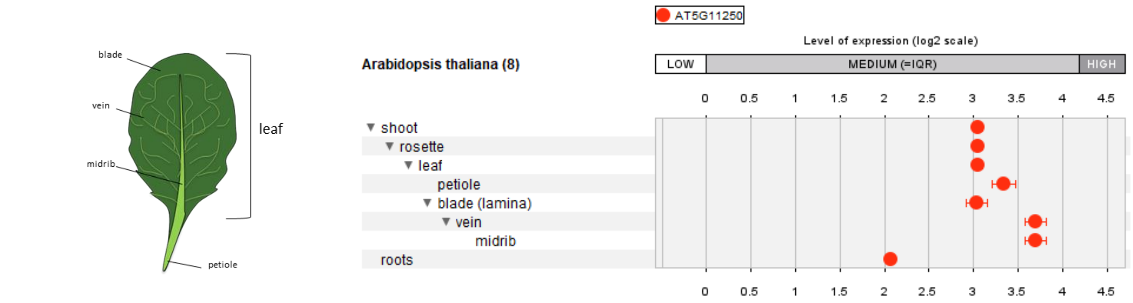


(b)

**Dataset:** 432 perturbations from data selection: AT_AFFY_ATH1-13

Showing 1 measure(s) of 1 gene(s) on selection:

Log2-ratio

-2.5 -2.0 -1.5 -1.0 -0.5 0.0 0.5 1.0 1.5 2.0 2.5

Down-regulated Up-regulated

83 of 432 perturbations fulfilled the filter criteria

Filter values for selected measure(s)

| Arabidopsis thaliana (83) | Pi-score | Log2-ratio ▼ | Fold-Change (\| 1.5 \|) | p-value (0.05) |
|---|---|---|---|---|
| AT-00202 (13) P. syringae pv. tomato study 10 (DC3000 hrpA) / P. syringae pv. tomat... | 7.17 | 2.65 | 6.14 | 0.002 |
| AT-00407 (24) Bay-0 parent / C24 [IAA study 9 (Bay-0) / IAA study 9 (C24)] | 5.69 | 1.77 | 3.37 | <0.001 |
| AT-00597 (3) EF-Tu (elf18) study 4 (Col-0) / mock treated seedling samples (Col-0) | 6.85 | 1.71 | 3.26 | <0.001 |
| AT-00407 (25) Bay-0 parent / C24 [untreated seedling samples (Bay-0) / untreated s... | 4.45 | 1.35 | 2.57 | <0.001 |
| AT-00407 (22) Bay-0 parent / C24 [IAA study 7 (Bay-0) / IAA study 7 (C24)] | 3.69 | 1.34 | 2.50 | 0.002 |
| AT-00407 (56) Sha / C24 [IAA study 9 (Sha) / IAA study 9 (C24)] | 3.71 | 1.33 | 2.48 | 0.002 |
| AT-00597 (11) Pep2 study 2 (Col-0) / mock treated seedling samples (Col-0) | 3.60 | 1.31 | 2.50 | 0.002 |
| AT-00597 (7) Pep2 (Col-0) / mock treated seedling samples (Col-0) | 4.59 | 1.31 | 2.47 | <0.001 |
| AT-00407 (89) Bl-1 / C24 [untreated seedling samples (Bl-1) / untreated seedling sa... | 5.17 | 1.29 | 2.45 | <0.001 |
| AT-00553 (3) H. arabidopsidis (Col-0; 6dpi) / untreated shoot samples (Col-0) | 1.74 | 1.24 | 2.32 | 0.039 |
| AT-00106 (16) P. syringae pv. tomato study 2 (DC3000 hrcC-) / P. syringae pv. tomat... | 3.80 | 1.19 | 2.28 | <0.001 |
| AT-00113 (16) salicylic acid / mock treated seedlings | 1.54 | 1.17 | 2.22 | 0.048 |
| AT-00253 (2) flg22 study 2 (3h) / H2O treated Col-0 seedlings (3h) | 3.07 | 1.17 | 2.25 | 0.002 |
| AT-00660 (5) IAA study 11 (3h) / mock treated root samples (3h) | 3.04 | 1.16 | 2.24 | 0.002 |
| AT-00407 (55) Sha / C24 [IAA study 8 (Sha) / IAA study 8 (C24)] | 4.00 | 1.12 | 2.17 | <0.001 |
| AT-00202 (10) P. syringae pv. tomato study 9 (DC3118 Cor-hrpS) / P. syringae pv. to... | 1.68 | 1.12 | 2.08 | 0.032 |
| AT-00106 (21) P. syringae pv. tomato study 3 (DC3000 hrcC-) / P. syringae pv. tomat... | 2.90 | 1.10 | 2.15 | 0.002 |
| AT-00413 (1) oxidative study 3 / Tween 20 treated aeral tissue samples | 1.50 | 1.09 | 2.11 | 0.043 |
| AT-00407 (57) Sha / C24 [untreated seedling samples (Sha) / untreated seedling sa... | 4.01 | 1.06 | 2.09 | <0.001 |
| AT-00407 (23) Bay-0 parent / C24 [IAA study 8 (Bay-0) / IAA study 8 (C24)] | 2.55 | 1.04 | 2.07 | 0.004 |
| AT-00407 (88) Bl-1 / C24 [IAA study 9 (Bl-1) / IAA study 9 (C24)] | 2.42 | 1.04 | 2.03 | 0.005 |
| AT-00407 (87) Bl-1 / C24 [IAA study 8 (Bl-1) / IAA study 8 (C24)] | 3.00 | 1.00 | 2.01 | 0.001 |
| AT-00543 (2) shift NPA to NAA (6h) / NPA study 3 | 3.21 | 0.99 | 1.98 | <0.001 |
| AT-00597 (1) EF-Tu (elf18) study 3 (Col-0) / mock treated seedling samples (Col-0) | 3.08 | 0.98 | 1.96 | <0.001 |
| AT-00407 (54) Sha / C24 [IAA study 7 (Sha) / IAA study 7 (C24)] | 1.98 | 0.97 | 1.94 | 0.009 |
| AT-00128 (4) EF-Tu (elf26) (60min) / untreated whole plant samples (Ler-0) | 2.09 | 0.93 | 1.90 | 0.006 |
| AT-00407 (34) Bay-0 parent / Fei-0 [IAA study 7 (Bay-0) / IAA study 7 (Fei-0)] | 2.67 | 0.86 | 1.81 | <0.001 |
| AT-00253 (1) flg22 study 2 (1h) / H2O treated Col-0 seedlings (1h) | 1.30 | 0.78 | 1.71 | 0.021 |
| AT-00494 (2) salicylic acid study 8 (Col-0) / mock treated leaf samples (Col-0) | 3.02 | 0.77 | 1.71 | <0.001 |
| AT-00693 (3) shift low to high light study 4 (Col-0) / low light study 5 (Col-0) | 1.31 | 0.74 | 1.66 | 0.017 |
| AT-00392 (2) flg22 study 3 (2h) / untreated leaf disc samples (Col-0) | 1.14 | 0.73 | 1.66 | 0.028 |
| AT-00407 (86) Bl-1 / C24 [IAA study 7 (Bl-1) / IAA study 7 (C24)] | 1.19 | 0.72 | 1.64 | 0.023 |
| AT-00407 (37) Bay-0 parent / Fei-0 [untreated seedling samples (Bay-0) / untreated s... | 1.55 | 0.72 | 1.65 | 0.007 |
| AT-00253 (3) OGs (1h) / H2O treated Col-0 seedlings (1h) | 1.25 | 0.72 | 1.64 | 0.018 |
| AT-00655 (6) IAA study 10 (12h) / mock treated root samples (12h) | 0.93 | 0.72 | 1.62 | 0.050 |
| AT-00276 (1) light study 6 (Col-0) / dark grown Col-0 seedlings | 1.55 | 0.71 | 1.63 | 0.007 |
| AT-00648 (2) P. cucumerina study 2 (Col-0) / mock inoculated rosette samples (Col-0) | 1.05 | 0.70 | 1.63 | 0.031 |
| AT-00638 (2) P. cucumerina (Col-0) / mock inoculated rosette samples (Col-0) | 1.21 | 0.69 | 1.62 | 0.018 |
| AT-00106 (19) P. syringae pv. tomato study 3 (DC3000 avrRpm1) / P. syringae pv. to... | 1.46 | 0.69 | 1.62 | 0.008 |
| AT-00128 (3) EF-Tu (elf26) (30min) / untreated whole plant samples (Ler-0) | 1.55 | 0.68 | 1.60 | 0.005 |
| AT-00593 (1) chitooctaose (Col-0) / mock treated whole plant samples (Col-0) | 2.42 | 0.67 | 1.59 | <0.001 |
| AT-00407 (101) Bl-1 / Fei-0 [untreated seedling samples (Bl-1) / untreated seedling ... | 2.07 | 0.66 | 1.58 | <0.001 |
| AT-00679 (1) RALF (30min) / mock treated seedling samples (30min) | 0.96 | 0.65 | 1.59 | 0.034 |
| AT-00407 (41) Bay-0 parent / Bur-0 [untreated seedling samples (Bay-0) / untreated ... | 1.04 | 0.62 | 1.54 | 0.021 |
| AT-00407 (40) Bay-0 parent / Bur-0 [IAA study 9 (Bay-0) / IAA study 9 (Bur-0)] | 1.78 | 0.62 | 1.54 | 0.001 |
| AT-00655 (4) IAA study 10 (4h) / mock treated root samples (4h) | 1.81 | 0.61 | 1.52 | 0.001 |
| AT-00655 (5) IAA study 10 (8h) / mock treated root samples (8h) | 1.08 | 0.60 | 1.51 | 0.016 |
| AT-00407 (13) IAA study 8 (Fei-0) / untreated seedling samples (Fei-0) | 1.10 | 0.58 | 1.51 | 0.013 |
| AT-00407 (70) Fei-0 / Col-0 [IAA study 7 (Fei-0) / IAA study 7 (Col-0)] | 1.22 | -0.59 | -1.51 | 0.009 |
| AT-00682 (2) low light + DBMIB (2h) / low light study 4 (6h) | 1.33 | -0.60 | -1.53 | 0.006 |
| AT-00407 (82) Bl-1 / Bay-0 parent [IAA study 7 (Bl-1) / IAA study 7 (Bay-0)] | 1.50 | -0.62 | -1.53 | 0.004 |
| AT-00407 (49) C24 / Fei-0 [untreated seedling samples (C24) / untreated seedling s... | 1.92 | -0.63 | -1.55 | <0.001 |
| AT-00106 (12) P. syringae pv. tomato study 2 (DC3000) / mock inoculated leaf sampl... | 1.34 | -0.65 | -1.56 | 0.009 |
| AT-00110 (2) ABA (3h) / mock treated seedlings (3h) | 0.93 | -0.66 | -1.59 | 0.040 |
| AT-00391 (20) flg22 study 10 (0h) / H2O pretreated leaf disc samples (Ler) | 1.07 | -0.67 | -1.57 | 0.025 |
| AT-00147 (1) B. cinerea / non-infected rosette leaf samples | 1.00 | -0.69 | -1.59 | 0.035 |
| AT-00650 (4) 24-eBL + glucose (dark) / mock treated seedling samples | 1.61 | -0.69 | -1.62 | 0.005 |
| AT-00391 (17) flg22 study 9 (0h) / H2O pretreated leaf disc samples (Ler) | 1.24 | -0.71 | -1.61 | 0.018 |
| AT-00681 (4) S. sclerotiorum study 2 (Col-0) / mock inoculated rosette leaf samples ... | 1.02 | -0.73 | -1.62 | 0.040 |
| AT-00407 (84) Bl-1 / Bay-0 parent [IAA study 9 (Bl-1) / IAA study 9 (Bay-0)] | 1.92 | -0.73 | -1.66 | 0.002 |
| AT-00407 (53) C24 / Bur-0 [untreated seedling samples (C24) / untreated seedling s... | 1.80 | -0.73 | -1.67 | 0.003 |
| AT-00650 (2) 24-eBL + glucose (dark) / 24-eBL (dark) | 2.42 | -0.75 | -1.68 | <0.001 |
| AT-00681 (2) S. sclerotiorum (Col-0) / mock inoculated rosette leaf samples (Col-0) | 1.83 | -0.78 | -1.73 | 0.004 |
| AT-00106 (13) P. syringae pv. tomato study 2 (DC3000 avrRpm1) / mock inoculated l... | 1.95 | -0.79 | -1.73 | 0.004 |
| AT-00407 (51) C24 / Bur-0 [IAA study 8 (C24) / IAA study 8 (Bur-0)] | 2.16 | -0.89 | -1.85 | 0.004 |
| AT-00407 (50) C24 / Bur-0 [IAA study 7 (C24) / IAA study 7 (Bur-0)] | 1.36 | -0.90 | -1.88 | 0.031 |
| AT-00682 (3) shift low to high light study 3 (0.5h) / low light study 4 (6h) | 2.51 | -0.91 | -1.89 | 0.002 |
| AT-00115 (2) zeatin study 3 (Col-0) / untreated whole plant samples (Col-0) | 2.91 | -0.97 | -1.97 | <0.001 |
| AT-00407 (47) C24 / Fei-0 [IAA study 8 (C24) / IAA study 8 (Fei-0)] | 2.53 | -0.99 | -1.99 | 0.003 |
| AT-00391 (21) flg22 study 11 (1h) / flg22 study 11 (0h) | 1.49 | -0.99 | -2.02 | 0.031 |
| AT-00682 (4) shift low to high light study 3 (2h) / low light study 4 (6h) | 2.81 | -1.03 | -2.04 | 0.002 |
| AT-00407 (42) C24 / Col-0 [IAA study 7 (C24) / IAA study 7 (Col-0)] | 2.41 | -1.07 | -2.08 | 0.006 |
| AT-00391 (23) flg22 study 11 (0h) / H2O pretreated leaf disc samples (Ler) | 2.16 | -1.13 | -2.13 | 0.012 |
| AT-00407 (52) C24 / Bur-0 [IAA study 9 (C24) / IAA study 9 (Bur-0)] | 3.02 | -1.15 | -2.19 | 0.002 |
| AT-00407 (45) C24 / Col-0 [untreated seedling samples (C24) / untreated seedling s... | 4.32 | -1.18 | -2.27 | <0.001 |
| AT-00407 (43) C24 / Col-0 [IAA study 8 (C24) / IAA study 8 (Col-0)] | 4.08 | -1.19 | -2.28 | <0.001 |
| AT-00106 (17) P. syringae pv. tomato study 3 (DC3000) / mock inoculated leaf sampl... | 3.71 | -1.23 | -2.35 | <0.001 |
| AT-00618 (5) high light study 7 (Col-0) / untreated leaf samples (Col-0) | 4.99 | -1.25 | -2.37 | <0.001 |
| AT-00541 (3) DFPM + ABA (Col-0) / solvent treated seedling samples (Col-0) | 2.11 | -1.29 | -2.38 | 0.023 |
| AT-00541 (1) ABA study 12 (Col-0) / solvent treated seedling samples (Col-0) | 3.62 | -1.37 | -2.56 | 0.002 |
| AT-00407 (44) C24 / Col-0 [IAA study 9 (C24) / IAA study 9 (Col-0)] | 3.72 | -1.47 | -2.77 | 0.003 |
| AT-00433 (7) ABA study 8 (Col-0) / solvent treated leaf samples (Col-0) | 4.72 | -1.81 | -3.52 | 0.002 |
| AT-00202 (11) P. syringae pv. tomato study 10 (DC3000) / mock inoculated leaf sam... | 6.01 | -2.19 | -4.50 | 0.002 |

(c)

**Dataset:** 189 perturbations from data selection: AT_mRNASeq_ARABI_GL-11

Showing 1 measure(s) of 1 gene(s) on selection:

Log2-ratio

-2.5 -2.0 -1.5 -1.0 -0.5 0.0 0.5 1.0 1.5 2.0 2.5

Down-regulated Up-regulated

20 of 189 perturbations fulfilled the filter criteria

Filter values for selected measure(s)

| Arabidopsis thaliana (20) | Pi-score | Log2-ratio ▼ | Fold-Change (\| 1.5 \|) | p-value (0.05) |
|---|---|---|---|---|
| AT-00795 (18) wounding (12h) / wounding (1h) | 8.07 | 2.77 | 6.86 | 0.001 |
| AT-00795 (16) wounding (6h) / wounding (1h) | 3.29 | 1.73 | 3.36 | 0.012 |
| AT-00795 (22) wounding (48h) / wounding (1h) | 3.38 | 1.51 | 2.87 | 0.006 |
| AT-00840 (11) shift dark to light (30min; cotyledon) / dark (cotyledon) | 3.29 | 1.26 | 2.40 | 0.002 |
| AT-00816 (13) GA3 (1h) / mock treated shoot samples (1h) | 4.78 | 1.20 | 2.29 | <0.001 |
| AT-00795 (20) wounding (24h) / wounding (1h) | 1.56 | 1.10 | 2.20 | 0.039 |
| AT-00795 (15) wounding (3h) / wounding (1h) | 2.38 | 1.05 | 2.07 | 0.005 |
| AT-00795 (19) wounding (12h) / wounding (6h) | 1.52 | 1.05 | 2.04 | 0.035 |
| AT-00840 (12) shift dark to light (30min; hypocotyl) / dark (hypocotyl) | 1.82 | 0.98 | 1.99 | 0.014 |
| AT-00816 (15) GA3 (4h) / mock treated shoot samples (4h) | 3.40 | 0.85 | 1.81 | <0.001 |
| AT-00840 (1) shift dark to light (1h; cotyledon) / dark (cotyledon) | 1.65 | 0.77 | 1.70 | 0.007 |
| AT-00836 (24) A. nodosum extract; P. cinnamomi (12hpi) / A. nodosum extract; P. cin... | 1.59 | 0.71 | 1.60 | 0.006 |
| AT-00840 (10) shift dark to light (15min; hypocotyl) / dark (hypocotyl) | 1.36 | 0.67 | 1.60 | 0.010 |
| AT-00754 (26) low P (24d) / untreated root samples (24d; sufficient P) | 1.10 | -0.58 | -1.51 | 0.013 |
| AT-00754 (6) C. incanum (24dpi; low P) / untreated root samples (24d; sufficient P) | 1.16 | -0.59 | -1.52 | 0.011 |
| AT-00840 (3) shift dark to light (3h; cotyledon) / dark (cotyledon) | 1.19 | -0.65 | -1.57 | 0.015 |
| AT-00812 (12) high light (24h) / high light (0.5h) | 1.56 | -0.71 | -1.64 | 0.006 |
| AT-00840 (5) shift dark to light (6h; cotyledon) / dark (cotyledon) | 1.31 | -0.82 | -1.76 | 0.025 |
| AT-00736 (1) B. cinerea (Col-0) / mock inoculated rosette leaf samples (Col-0) | 2.29 | -0.86 | -1.82 | 0.002 |
| AT-00795 (21) wounding (24h) / wounding (12h) | 2.89 | -1.67 | -3.12 | 0.019 |

**Figure S4.** Expression data for BNT1 (AT5G11250) were obtained using the GENEVESTIGATOR® Anatomy (a) or Perturbations (b) tools for Arabidopsis datasets.

(a) The different tissues have been arranged into logical hierarchies. Points indicate mean values, and whiskers indicate standard errors.

(b,c) The tables list experimental conditions that lead to up-or down-regulation of BNT1. Perturbations are ordered by the highest fold change at the top. Only comparisons with a fold-change of at least 1.5 and a p-value ≤ 0.05 are included. The Log2 ratio indicates the magnitude of the difference between the average expression in biotic stress samples and the average expression in control samples.
