## Supplementary material for "The *Arabidopsis* TNL immune receptor BNT1 localizes to the plastid envelope and mediates flg22-induced resistance against *Pseudomonas*": Figure S5

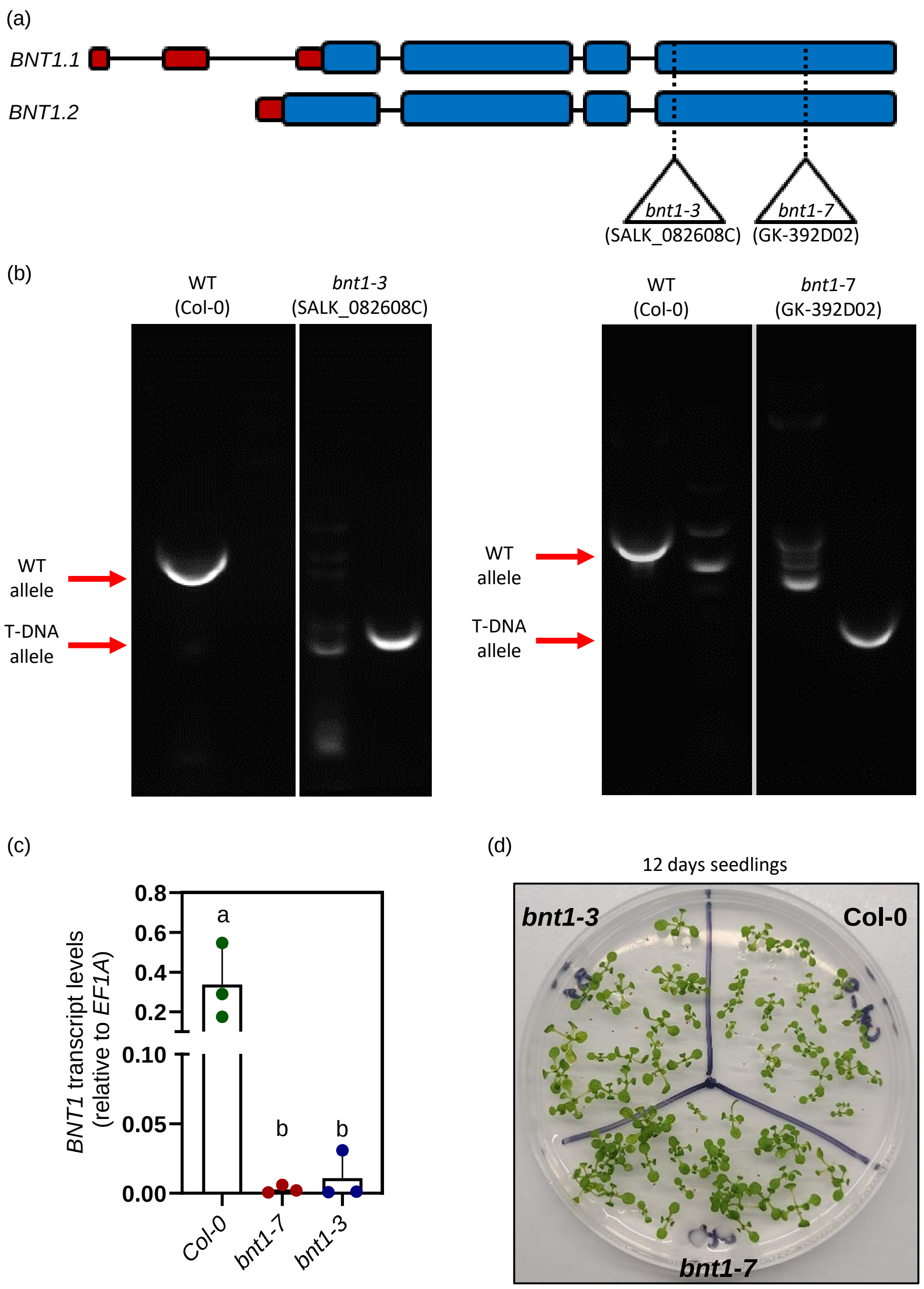


**Figure S4.** Characterization of T-DNA insertion mutant lines.

(a) Schematic representation of T-DNA insertion sites in the *BNT1.1* and *BNT1.2* gene models in *Arabidopsis*. Dashed lines and triangles: T-DNA insertion sites of mutant allele lines used (*bnt1-3*, SALK_082608C and *bnt1-7*, GK-392D02). Blue blocks: coding exons, black lines: introns, red blocks: 5'UTR exons.

(b) Genotypification PCR confirming WT Col-0 and T-DNA mutant allele in the lines used (*bnt1-3*, SALK_082608C; *bnt1-7*, GK-392D02).

(c) Relative transcript levels of total *BNT1* quantified by RT-qPCR in *Arabidopsis* WT Col-0 and T-DNA mutant allele lines used (*bnt1-3*, SALK_082608C; *bnt1-7*, GK-392D02), using the specific primers indicated in Table S1. The mean ± SE from three independent experiments is shown (each data point with 3 leaves from 3 different plants pooled together for RNA extraction). Individual data points are presented as scatter-dots. Different letters indicate statistically significant differences ($p < 0.05$, ANOVA, Tukey's multiple comparison test).

(d) Representative picture of 12-day-old WT Col-0, *bnt1-3* and *bnt1-7* seedlings grown on horizontal ½ MS agar plates.
