## Supplementary material for "The *Arabidopsis* TNL immune receptor BNT1 localizes to the plastid envelope and mediates flg22-induced resistance against *Pseudomonas*": Figure S6

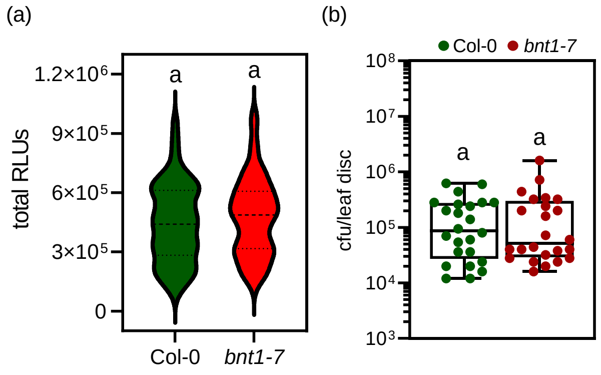


**Figure S6.** Total apoplastic reactive oxygen species (ROS) and growth of sprayed *Pst* in WT (Col-0) and *bnt1-7* mutant plants.

(a) ROS production in leaf discs was quantified after treatment with mock or 100 nM flg22. Relative light units (RLUs) from all time points were integrated over 30 minutes for each leaf disc. Data from three independent experiments are presented as violin plots (at least 22 biological replicates per experiment; n = 66 discs). Different letters indicate statistically significant differences ($p < 0.05$, Student's *t* test).

(b) Growth of the virulent bacteria *Pst* 5 days post-infection. *Pst* was sprayed ($OD_{600}$=0.01). The number of cfu per leaf disc is presented as scatter-dots in the boxplots. The data from three independent experiments is shown (each one with 8 biological replicates; n=24). Different letters indicate statistically significant differences between genotypes ($p < 0.05$, analysis of variance (ANOVA), Fisher's LSD test).
