## Supplementary material for "The *Arabidopsis* TNL immune receptor BNT1 localizes to the plastid envelope and mediates flg22-induced resistance against *Pseudomonas*": Figure S7

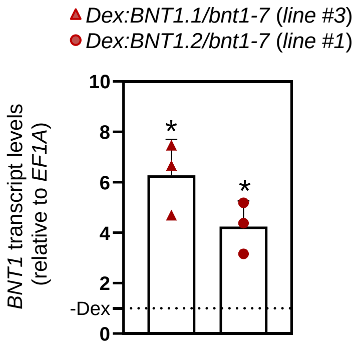


**Figure S7.** Characterization of *Arabidopsis* transgenics lines Dex:BNT1.1-GFP/*bnt1-7* (*line #3*) and Dex:BNT1.2:GFP/*bnt1-7* (*line #1*).
Relative transcript levels of total *BNT1* quantified by RT-qPCR in *Arabidopsis* leaves infiltrated with or without 30 µM dexamethasone (Dex). The relative average total *BNT1* levels without Dex treatment (-Dex) is shown as a dashed line in the graph. The mean ± SE from three independent experiments is shown (each data point with 3 leaves from 3 different plants of same line pooled together for RNA extraction). Individual data points are presented as scatter-dots. Asterisks indicate statistically significant differences compared to -Dex treatment ($p < 0.05$, Student’s *t* test).
