## Supplementary material for "The *Arabidopsis* TNL immune receptor BNT1 localizes to the plastid envelope and mediates flg22-induced resistance against *Pseudomonas*": Figure S8

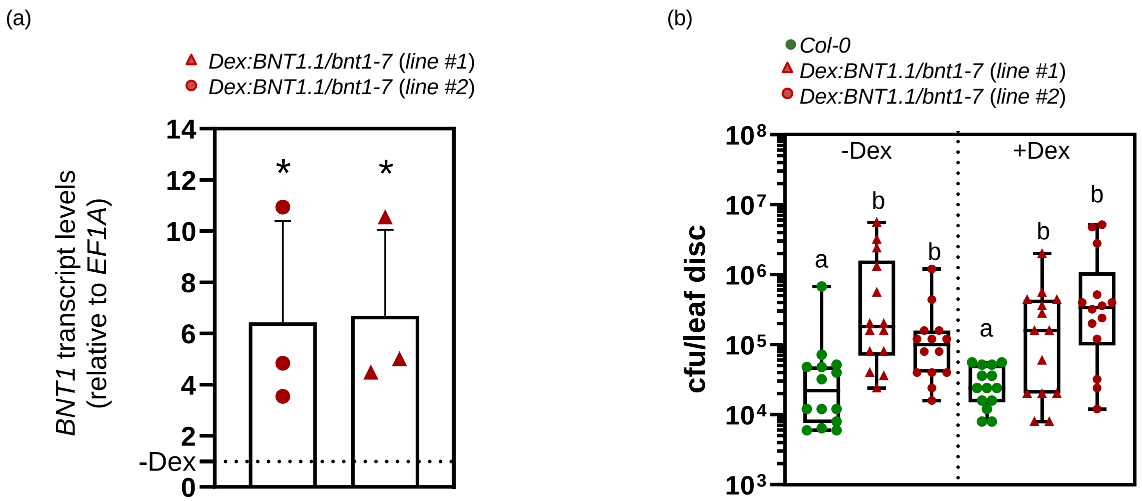


**Figure S8.** Characterization of *Arabidopsis* transgenics lines Dex:BNT1.1-GFP/*bnt1-7* (lines #*1* and #*2*).

(a) Relative transcript levels of total *BNT1* quantified by RT-qPCR in *Arabidopsis* leaves infiltrated with or without 30 µM dexamethasone (Dex). The relative average *BNT1* level without Dex treatment (-Dex) is shown as a dashed line in the graph. The mean ± SE from three independent experiments is shown (each data point with 3 leaves from 3 different plants of same line pooled together for RNA extraction). Individual data points are presented as scatter-dots. Asterisks indicate statistically significant differences compared to -Dex treatment ($p < 0.05$, Student's *t* test).

(b) As shown in Figure 6b, WT Col-0 or transgenic plant lines Dex:*BNT1.1*-GFP/*bnt1-7* were infiltrated with 30 µM Dex to induce *BNT1.1* expression in *bnt1* background. After 24 hours, the plants were treated with 1 µM flg22. *Pst* was inoculated into the same leaf one day later (OD=0.005), and bacterial growth was measured three days post-infection. Number of colony-forming units (cfu) per leaf disc is presented as scatter-dots in the boxplots. The mean ± SE from two independent experiments is shown (each one with 7 biological replicates; n = 14). Different letters indicate statistically significant differences between genotypes with or without Dex ($p < 0.05$, ANOVA, Fisher's LSD test).
