## Supplementary material for "The *Arabidopsis* TNL immune receptor BNT1 localizes to the plastid envelope and mediates flg22-induced resistance against *Pseudomonas*": Table S1

**Table S1.** Vectors, constructs and primers list used in this study

| Plasmid/constructs | Description | Antibiotic^(a)^ | Reference |
| --- | --- | --- | --- |
| pBAV150 | Gateway binary plant expression vector (*Dex* promoter, C-terminal GFP-tag) | Km^R^/BASTA^R^ | Vinatzer et. al 2006 |
| pGWB405 | Gateway binary plant expression vector (*CaMV:35S* promoter, C-terminal GFP-tag) | Sp^R^/Km^R^ | Nakagawa et. al 2007 |
| pGWB405-I BNT1.2^Δ31-1190^-GFP | CaMV 35S:BNT1 variant:GFP in a binary vector | Sp^R^/Km^R^ | This study |
| pGWB405-II- BNT1.2^Δ34-1190^-GFP | CaMV 35S:BNT1 variant:GFP in a binary vector | Sp^R^/Km^R^ | This study |
| pGWB405-III- BNT1.2^Δ43-1190^-GFP | CaMV 35S:BNT1 variant:GFP in a binary vector | Sp^R^/Km^R^ | This study |
| pGWB405-IV- BNT1.2^Δ57-1190^-GFP | CaMV 35S:BNT1 variant:GFP in a binary vector | Sp^R^/Km^R^ | This study |
| pGWB405-BNT1.1^Δ158-1107^-GFP | CaMV 35S:BNT1.1TIR:GFP in a binary vector | Sp^R^/Km^R^ | This study |
| pGWB405-BNT1.2^Δ240-1189^-GFP | CaMV 35S:BNT1.2TIR:GFP in a binary vector | Sp^R^/Km^R^ | This study |
| pBAV150-BNT1.1:GFP | Dex:BNT1.1:GFP in a binary vector | Km^R^ /BASTA^R^ | This study |
| pBAV150-BNT1.2:GFP | Dex::BNT1.2:GFP in a binary vector | Km^R^ /BASTA^R^ | This study |
| pBAV154- OEP7:RFP | Dex:OEP7:RFP:HA in a binary vector | Km^R^ /BASTA^R^ | Cecchini et. al 2015 |
| pBAV154- BiP:RFP | Dex:BiP:RFP:HA in a binary vector | Km^R^ /BASTA^R^ | Cecchini et. al 2015 |

(a) BASTA^R^, BASTA (glufosinate ammonium) resistance; Km^R^, kanamycin resistance; Sp^R^, spectinomycin resistance.

| Name Primers | | Sequence (5’-3’) | Purpose |
| --- | --- | --- | --- |
| ERF1alpha qPCR FW | | GAGCGGGAAATTGTCAGGGA | RT-qPCR (Figure 5c, Figure 3c-e, Figure 6a, Figure S5c, Figure S7 and Figure S8a) |
| ERF1alpha qPCR RV | | GAAACGCTCAGCACCAAT | RT-qPCR (Figure 5c, Figure 3c-e, Figure 6a, Figure S5c, Figure S7 and Figure S8a) |
| YLS8 qPCR FW | | AAGATCAACTGGGCTCTCAAGG | RT-qPCR (Figure 3d and Figure S3b,c) |
| YLS8 qPCR RV | | TGGGAAGCTCGATTAGTAACGG | RT-qPCR (Figure 3d and Figure S3b,c) |
| BNT1.1-cacc-FW | caccATGGAGTTTCAAAGAATGGGAATCA | | BNT1.1 full and BNT1.1 TIR cloning for constructs |
| BNT1.2-cacc-FW | caccATGGCTTCTTCGTTTTTCCTTACCAC | | BNT1.2 full and variants (I to IV) cloning for constructs |
| BNT1-nostop-RV | GAATGCGGGTTACGCCAACTATCG | | cloning for constructs |
| BNT1.2^Δ31-1190-^RV | GATTCCATCATGATAACAAA | | I BNT1.2 variant cloning for constructs |
| BNT1.2^Δ34-1190-^RV | TGATAACAAAGAAAAGAAG | | II BNT1.2 variant cloning for constructs |
| BNT1.2^Δ43-1190-^RV | CTTTATCTCCTTCATCATCt | | III BNT1.2 variant cloning for constructs |
| BNT1.2^Δ57-1190-^RV | CGAAGAAGGTGGAGGAACTG | | IV. BNT1.2 variant cloning for constructs |
| BNT1.2^Δ240-1189-^FW | caccatgACACACCAAGTCTTTCCCAG | | BNT1.2 TIR cloning for constructs |
| BNT1-TIRnostop-RV | AGTGGAATTAATCAATATGT | | BNT1 variants (I to IV) (TIR1.1-TIR1.2) cloning for constructs |
| PR1_FW | GTAGGTGCTCTTGTTCTTCCC | | RT-qPCR (Figure 5c) |
| PR1_RV | CACATAATTCCCACGAGGATC | | RT-qPCR (Figure 5c) |
| BNT1.1_FW | GAGATGAGCAATCATCTTCCGC | | RT-qPCR (Figure 3c-e and Figure 6a) |
| BNT1.2_FW | ATGGCTTCTTCGTTTTTCCTTACC | | RT-qPCR (Figure 3c-e and Figure 6a) |
| BNT1/1.1/1.2 RV | CTTCATTATCCCAGTTGATTGAATGG | | RT-qPCR (Figure 3c-e, Figure 6a and Figure S5c) |
| GC1_FW | TCGTCCAAGAATCAATTGTGGGC | | RT-qPCR (Figure S3b) |
| GC1_RV | GTGTTGCCGGAGGTTCCCGG | | RT-qPCR (Figure S3b) |
| LHCB2.1FW | TTGGTGTATCCGGTGGTGGCC | | RT-qPCR (Figure S3b) |
| LHCB2.1RV | GTCCGTACCAGATGCTTTGAGGAGTAGA | | RT-qPCR (Figure S3b) |
| SULTR2.1FW | GGTGTTGAGCTAGTGATCGTTAACCCG | | RT-qPCR (Figure S3b) |
| SULTR2.1RV | CCCGTAACACAACTGGTCCTTTGA | | RT-qPCR (Figure S3b) |
| ATTB2 RV | ACCACTTTGTACAAGAAAGCTGGGT | | RT-qPCR (Figure S7 and Figure S8a) |
| BNT1.C-TERMFW | GAGAGGCCTCTTCCTACATC | | RT-qPCR (Figure S5c, Figure S7 and Figure S8a) |
